# Assessing the impact of whole genome duplication on gene expression and regulation during arachnid development

**DOI:** 10.1101/2024.12.18.628675

**Authors:** Madeleine E. Aase-Remedios, Daniel J. Leite, Ralf Janssen, Alistair P. McGregor

**Affiliations:** Department of Biosciences, Durham University, Durham, DH1 3LE, UK; Department of Earth Sciences, Uppsala University, 752 36, Uppsala, Sweden

**Keywords:** whole genome duplication, evolution, arachnids, spiders, gene regulation

## Abstract

Whole genome duplication (WGD) generates new genetic material that can contribute to the evolution of the regulation of developmental processes and phenotypic diversification. A WGD occurred in an ancestor of arachnopulmonates (spiders, scorpions, and their relatives), which provides an important independent comparison to WGDs in other animal lineages like those in vertebrates and horseshoe crabs. After WGD, arachnopulmonates retained many duplicated copies (ohnologues) of developmental genes including clusters of homeobox genes, many of which have been inferred to have undergone subfunctionalisation. However, there has been little systematic analysis of gene regulatory sequences and comparison of the expression of ohnologues versus their single-copy orthologues between arachnids. Here we compare the regions of accessible chromatin and gene expression of ohnologues and single-copy genes during three embryonic stages between an arachnopulmonate arachnid, the spider *Parasteatoda tepidariorum*, and a non-arachnopulmonate arachnid, the harvestman *Phalangium opilio*. We found that the expression of each spider ohnologue was lower than their single-copy orthologues in the harvestman providing evidence for subfunctionalisation. However, this was not reflected in a reduction in peaks of accessible chromatin because both spider ohnologues and single-copy genes had more peaks than the harvestman genes. We also found peaks of open chromatin that increased in the late stage associated with activation of genes expressed later during embryogenesis in both species. Taken together, our study provides the first genome-wide comparison of gene regulatory sequences and gene expression in arachnids and thus provides new insights into the impact of the arachnopulmonate WGD.

**Significance statement:** The comparison of independent WGD events is essential to understanding their evolutionary outcomes. Here we examine gene expression and chromatin accessibility during the development of two arachnids, one of which underwent an ancient WGD. Our data provide the first direct comparison of the regulation of ohnologues and single-copy genes in arachnids.

## Introduction

Whole genome duplication (WGD) generates new genetic material that can contribute to evolutionary diversification (Sémon and Wolfe 2007; Van de Peer et al. 2017; MacKintosh and Ferrier 2018). WGDs have occurred several times during animal evolution including the two rounds (2R) on the vertebrate stem, three in horseshoe crabs and a single event in the ancestor of arachnopulmonates, a lineage of arachnid arthropods including spiders, scorpions, and their relatives (Putnam et al. 2008; Kenny et al. 2016; Schwager et al. 2017; Nong et al. 2021; Kulkarni et al. 2024; Marlétaz et al. 2024) (Figure 1). While the vertebrate 2R WGD have been intensively studied, fully understanding the consequences of WGDs requires comparison between independent events.

**Figure 1:**
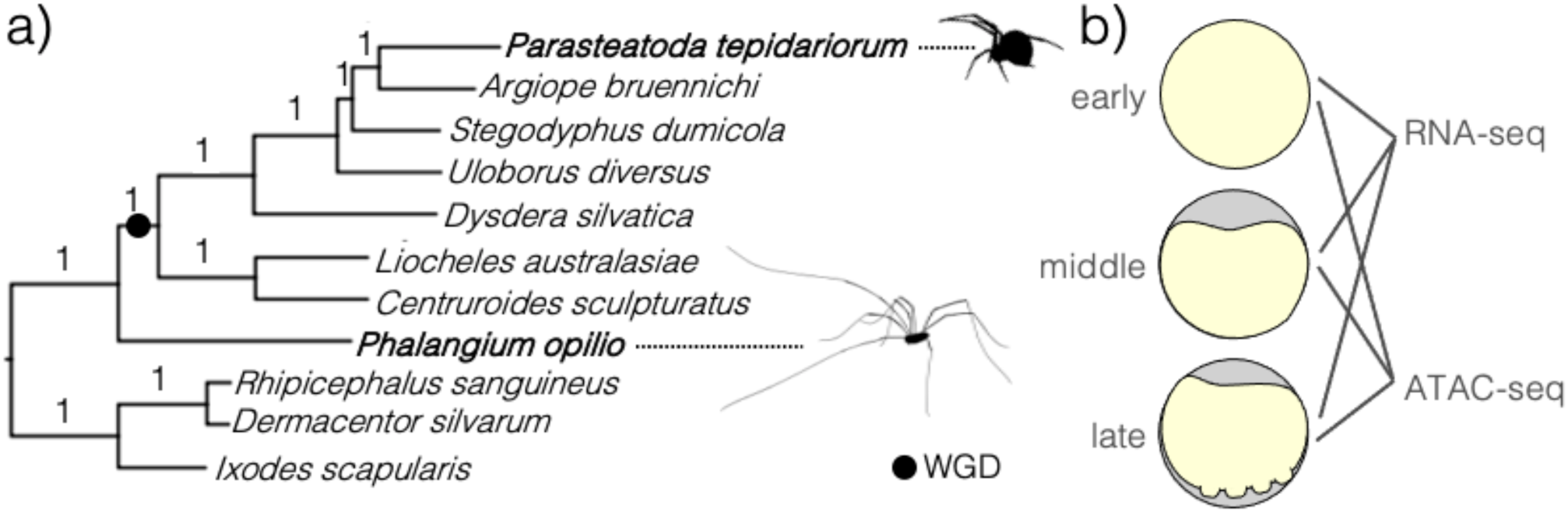
a) The *OrthoFinder* tree of orthogroups (OGs) with one gene in all species for the 11 arachnid proteomes used in this study. The two focal species, *P. tepidariorum* and *P. opilio*, are in bold text. Branches are labelled with STAG support proportions. Silhouettes were taken from Phylopic.org. b) Schematic of early, middle, and late stages used for ATAC and RNA sequencing, corresponding to *P. tepidariorum* and *P. opilio* stages 5, 7, and 8 (Mittmann and Wolff 2012; Gainett et al. 2022).

Previous work has shown that the arachnopulmonate WGD resulted in spiders, scorpions, pseudoscorpions, amblypygids and uropygids having retained WGD paralogues (ohnologues) of many key developmental genes, for example, homeobox genes (including two Hox gene clusters), Sox genes, micro-RNAs, retinal determination genes, and Wnt-, Hedgehog- and Notch-Delta-signalling components, compared to other arachnid lineages that did not experience this event such as harvestmen, which have only one copy (Figure 1) (Schwager et al. 2007; Leite et al. 2016; Schwager et al. 2017; Leite et al. 2018; Akiyama-Oda and Oda 2020; Baudouin-Gonzalez et al. 2021; Janssen et al. 2021; Aase-Remedios et al. 2023; Baudouin-Gonzalez et al. 2023; Kulkarni et al. 2024). This pattern is similar to the repertoire of retained ohnologues of vertebrates relative to pre-duplicate genes in chordates like amphioxus or tunicates (Dehal 2002; Mazet and Shimeld 2002; Cañestro et al. 2003; Woolfe and Elgar 2007; Putnam et al. 2008; Holland 2013).

Developmental genes involved in transcriptional regulation and signalling make up a larger proportion of ohnologues than expected by chance in vertebrates (Mazet and Shimeld 2002). The sub- and neo-functionalisation of these ohnologues may constitute the mechanism by which WGDs can facilitate the diversification of developmental processes, and potentially evolutionary innovations (Cañestro et al. 2013; Mottes et al. 2021). Comparisons of gene expression and chromatin profiling between 2R vertebrates and the pre-duplicate state represented by the invertebrate chordate amphioxus revealed widespread subfunctionalisation, as well as a different fate for widely-expressed genes where one ohnologue underwent specialisation (Marlétaz et al. 2018). While the primary model of subfunctionalisation posits that lower expression between ohnologues occur by mutations disrupting and degrading *cis*-regulatory elements (Force et al. 1999), the specialised vertebrate ohnologues were expressed in a highly restricted pattern relative to the ancestral gene but gained *cis*-regulatory elements consistent with a fine-tuning of regulatory complexity (Marlétaz et al. 2018). Both these mechanisms may contribute to the macroevolutionary consequences of WGD, however, regulatory data from lineages besides vertebrates is scarce. We have no knowledge of how the arachnopulmonate WGD has impacted the gene regulation in spider genomes during development, although there is evidence of subfunctionalisation of some developmental genes, including homeobox genes, during spider embryogenesis (Janssen et al. 2015; Leite et al. 2018; Baudouin-Gonzalez et al. 2021; Aase-Remedios et al. 2023; Janssen and Pechmann 2023).

To investigate this further, we carried out RNA-sequencing and characterised accessible chromatin using assay for transposase-accessible chromatin using sequencing (ATAC-seq) in the spider *Parasteatoda tepidariorum* and the harvestman *Phalangium opilio* (Figure 1). For each species we investigated three morphologically comparable developmental stages encompassing the formation of the germ band and the onset of segmentation when many key developmental genes are expressed and patterning is initiated by the Hox genes (Mittmann and Wolff 2012; Gainett et al. 2022). We also sequenced a contiguous genome assembly for the harvestman to serve as an improved reference for our data and better enable comparisons among arachnids, as the Arachnopulmonata-Opiliones ancestor is more recent than the ancestor between arachnopulmonates and the other arachnids with reference genomes like ticks or mites (Figure 1).

With these data, we found that the spider *P. tepidariorum* has a larger genome with longer intergenic and intronic distances, which contains more peaks of accessible chromatin than the harvestman *P. opilio*. Moreover, spider ohnologues belong to gene families with longer intergenic distances and longer introns and have more peaks of accessible chromatin than the genes in single copy families. These characteristics are true even of the harvestman pro-orthologues of spider ohnologues and orthologues of spider single-copy genes. We also examined the regulatory states for genes that were activated in the late stage of our experiment (Figure 1B), identifying peaks associated with their expression patterns. We found instances of conserved 1:2 synteny for a conserved late-specific locus and among homeobox gene clusters, consistent with WGD. These data provide the first glimpse into the impact of the arachnopulmonate WGD on *cis*-regulatory sequences across the genome and gene expression at key developmental stages.

## Methods

### RNA-seq, ATAC-seq protocols

We aimed to compare the RNA expression and chromatin accessibility during development between a spider and a comparable arachnid that had not undergone the arachnopulmonate WGD. We sampled embryos of the harvestman *P. opilio* collected from a wild population in Uppsala, Sweden in October 2023. Due to somewhat non-synchronous development, batches of embryos were dechorionated with 1:1 bleach:water solution and sorted into pre-germ band (stage 5), germ-band (stage 7), and limb-bud/onset of segmentation (stage 8) stages (Gainett et al. 2022) (Figure 1B). Samples of 25, 15, and 10 embryos for each stage, respectively, were collected for ATAC protocols (Diagenode C01080002) to account for fewer cells in younger embryos, and the remaining embryos from each batch were used for RNA extraction with TRIzol in four replicates for each stage. The standard Diagenode protocol was carried out four times for each stage, with the additional step of using a haemocytometer to determine the amount of lysate containing 50,000 nuclei to use for tagmentation. We then sampled equivalent stages of embryos of *P. tepidariorum* from the lab culture in Durham, UK, for ATAC sequencing in four replicates for each stage, and conducted the same protocol as above. We sampled three replicates for the early (st5) and middle (st7) stages, and four replicates for the late (st8) stage of *P. tepidariorum* embryos for RNA extraction (Mittmann and Wolff 2012). ATAC libraries for the twelve samples of each species were pooled at equimolar concentrations for sequencing.

### High Molecular Weight (HMW) DNA extraction

We aimed to improve the reference genome assembly of the harvestman *P. opilio* with long-read sequencing. We sampled the anterior part of the body up to the fourth limb-bearing segment and including the limbs of a single female *P. opilio* caught from the wild population in Uppsala, Sweden. We extracted HMW DNA with the Monarch kit (T3060S), but instead of using the beads to precipitate DNA, we used a centrifuge (5000 rpm, 15 minutes) to pellet the DNA after the addition of isopropanol, then continued with the washing steps.

### Quality control and sequencing

ATAC libraries were quality checked on the Agilent TapeStation with a D1000 high-sensitivity assay. The HMW DNA extraction concentration was measured with a Qubit spectrophotometer and quality checked on the Agilent TapeStation with the Genomic DNA assay. ATAC libraries were paired-end sequenced (150 bp) on an Illumina NovaSeq by Novogene, yielding 2.32 B (407 Gb) and 2.71 B (347 Gb) reads for *P. opilio* and *P. tepidariorum*, respectively. cDNA libraries were prepared from RNA extractions with poly-A enrichment and paired-end sequenced with 150 bp fragments as well, yielding a total of 1.18 B reads (177.3 Gb) for both species, with an average of 54 million reads per sample.

Libraries for the HMW DNA were prepared for PacBio Revio Hi-Fi sequencing by Novogene, yielding 8.2 M reads (101.11 Gb).

### Draft harvestman genome assembly

Adapters were removed with *HiFiAdapterFilt* (Sim et al. 2022), then the mitogenome was assembled with *MitoHiFi v2.2* (Uliano-Silva et al. 2023) and reads not mapping to it were isolated with *minimap2 v2.24* (Li 2021). These non-mitochondrial reads were assembled with *hifiasm v0.19.9* with the *s* parameter set to 0.2 for higher heterozygosity due to the sample originating from a wild population (Cheng et al. 2021), and the assembly was reduced to a single pseudohaplotype with *purge_dups v1.2.6* (Guan et al. 2020). The assembly was then aligned with *blastn* (Altschul et al. 1990) to the *Genbank Refseq* RNA database (ftp://ftp.ncbi.nlm.nih.gov/blast/db/refseq_rna.14.tar.gz) to identify contaminants. Read coverage was determined with *minimap2* (Li 2021) and completeness was calculated with *BUSCO v5.5.0* using the *arachnida_odb10* reference (Manni et al. 2021). These genome statistics were visualised with *BlobTools2* and the assembly was filtered to remove sequences with GC proportions below 0.35 or above 0.45, coverage lower than 20, length shorter than 10 kb, and hits from non-animal kingdoms (Challis et al. 2020). The final assembly comprises 649 Mb across 123 contigs, with an N50 of 9.88 Mb and a BUSCO completeness score of 93.6%. Repeats were identified with *RepeatModeler* with *LRTStruct* and *quick* options and masked with *RepeatMasker* with *xsmall*, *gff*, and *rmblast* options (Smit and Hubley 2023). Assembly statistics can be found in Figure S1. Raw HiFi reads were deposited in GenBank with the identifier SRR31693326 under BioProject PRJNA1197178 and will be released upon publication. This final assembly was deposited in GenBank under the accession JBJXVG000000000. The version described in this paper is version JBJXVG010000000.

### RNA-seq analysis

To compare RNA expression between *P. opilio* and *P. tepidariorum* we generated gene annotations using RNA-seq evidence for in silico prediction with *BRAKER3* (Gabriel et al. 2024). The RNA-seq data generated in this study for *P. opilio* and RNA sequencing reads from pooled developmental stages of *P. tepidariorum* (GenBank: SRR1507193 and SRR1507194) were trimmed of adapters and low-quality bases using *cutadapt v4.1* (Martin 2011). Trimmed reads were aligned to the reference genome of *P. tepidariorum* (Zhu et al. 2023) with our annotation or the *de novo* assembly and annotation for *P. opilio* with *STAR v2.7.9a* (Dobin et al. 2013). We used *DESeq2* in *R v4.3.1* for differential gene expression analysis (Love et al. 2014). To enable comparisons between sequencing runs and between species, we standardised the number of reads across samples and between species, based on the number of mapped reads per sample per species. Raw RNA-seq read were deposited in GenBank under the BioProject PRJNA1198402 and will be released upon publication.

### ATAC-seq analysis

To compare chromatin accessibility between *P. opilio* and *P. tepidariorum* we called ATAC-seq peaks for our three developmental stages. Adapters were removed with *cutadapt v4.1* as above and reads were aligned to the reference genomes with *Bowtie2 v2.4.5* (Langmead and Salzberg 2012). Using *SAMtools v1.17* (Danecek et al. 2021), alignments were filtered, sorted, duplicate reads were removed, and replicate runs of samples were merged. We recentred reads as previously (Buenrostro et al. 2015) and calculated effective genome size with *khmer v2.1.2* (Crusoe et al. 2015) before peak calling with *MACS2 v2.2.7.1* (Zhang et al. 2008). We used *DiffBind* in *R v4.3.1* to determine significant differences in peaks between time points (Stark and Brown 2011). For each gene for each developmental stage, we also tallied the number of peaks between each gene and its up- and down-stream neighbour as well as those within the introns of the gene, and the total height of the peak(s) overlapping the region directly adjacent to its transcription start site (TSS). For the spider, this was the start position ± 500 bp, and for the harvestman, this was the start position ± 300 bp, to scale for differences in genome size. Raw ATAC reads were deposited in GenBank under the BioProject PRJNA1199734 and will be released upon publication.

### Ohnologue definition

We used *OrthoFinder* (Emms and Kelly 2019) to define orthogroups (OGs) in 11 available arachnid proteomes, using only the longest isoform for each gene (Table 1). To find ohnologues for our species comparison, these OGs were filtered to those with an average number of genes among WGD species between 1.5 and 2.5, an average number of genes between 0.5 and 1.5 among non-WGD species, one copy in *P. opilio* and two copies in *P. tepidariorum*, and the two spider genes in the OG had to be on different chromosomes in at least two out of the three spiders with chromosome-level assemblies in our analysis (*P. tepidariorum*, *Argiope bruennichi*, and *Dysdera silvatica*). This resulted in 530 ohnologue OGs. Single-copy orthologues were defined by *OrthoFinder* as the OGs containing a single gene across all 11 species (n=1,435). Within ohnologous OGs, the A ohnologue was defined as the higher expressed ohnologue, and the B as the lower, unless ohnologues had been previously designated A or B in publications.

**Table 1:** Species and genome accessions used in OrthoFinder. The left-most column indicates the species that share the arachnopulmonate WGD. The *P. opilio* genome can be accessed in our repository.

|  | Species | Accession |
| --- | --- | --- |
| WGD | *Parasteatoda tepidariorum* | Science Data bank doi:10.11922/sciencedb.o00019.00014 |
| WGD | *Argiope bruennichi* | GenBank: GCA_947563725.1 |
| WGD | *Stegodyphus dumicola* | GenBank: GCA_010614865.2 |
| WGD | *Uloborus diversus* | GenBank: GCA_026930045.1 |
| WGD | *Dysdera silvatica* | https://github.com/molevol-ub/Dysdera_silvatica_genome |
| WGD | *Liocheles australasiae* | So et al. 2023 |
| WGD | *Centruroides sculpturatus* | GenBank: GCA_000671375.2 |
|  | *Phalangium opilio* | This study: JBJXVG000000000 |
|  | *Rhipicephalus sanguineus* | GenBank: GCA_013339695.2 |
|  | *Dermacentor silvarum* | GenBank: GCA_013339745.2 |
|  | *Ixodes scapularis* | GenBank: GCA_016920785.2 |

### Functional annotation

To functionally characterise *P. opilio* and *P. tepidariorum* proteomes, we ran *InterProScan v5.70-102.0* and used the corresponding Gene Ontology (GO) lookup table to find GO terms associated with each protein (Mitchell et al. 2015; Blum et al. 2021) for submission to *GO-Compass* (Harbig et al. 2023).

## Results

### Expression and chromatin dynamics across stages

We first compared the expression and chromatin accessibility among 530 ohnologue gene families and 1,435 single copy orthologue gene families between the three developmental stages of the spider *P. tepidariorum* and the harvestman *P. opilio* (Figure 2a). We found that expression levels were largely consistent between stages for both ohnologue and single-copy orthologue gene families, except for a slightly significant decrease in expression of harvestman ohnologue family pro-orthologues between early and late stages (Figure 2c).

**Figure 2:**
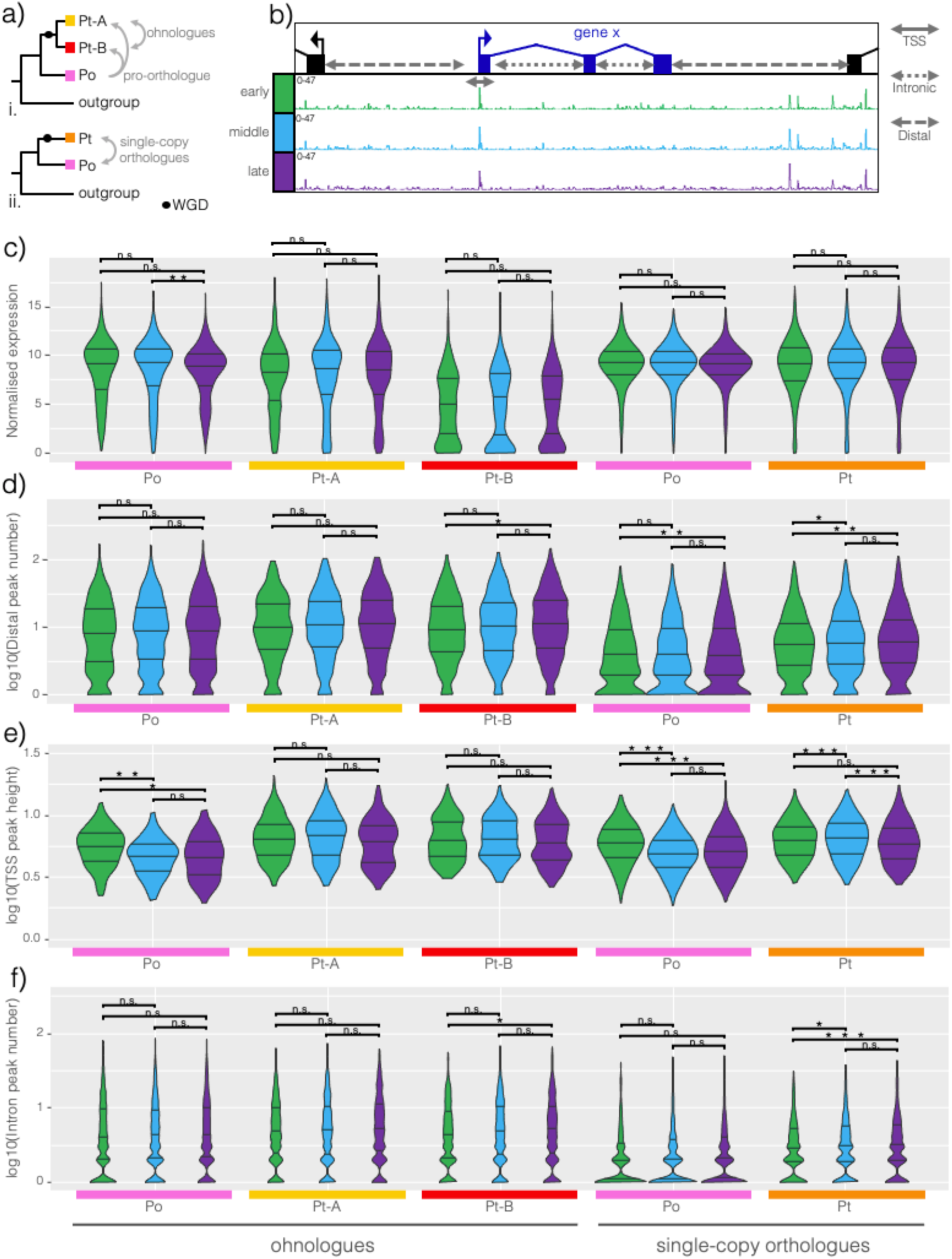
Expression and peak dynamics between stages for ohnologue families and single-copy orthologue families. a) Schematic illustrating the relationships between harvestman and spider genes in the case i. where both ohnologues are retained or ii. where one is lost and the spider retains a single orthologue. b) Schematic defining the intronic, distal, and transcript start site (TSS) regions assigned to each gene with example ATAC profiles for each stage. c) – f) Violin plots of normalised reads, number of peaks in the distal region, height of peaks overlapping with the TSS, and number of peaks within the introns for harvestman and spider ohnologue gene families and single-copy orthologue gene families. Significance levels were determined with a two-tailed Wilcoxon rank sum test (*** p<0.001; ** p<0.01; * p<0.05; n.s. p>0.05).

We then examined the number of distal peaks annotated to each gene’s up- and down-stream region, TSS peaks annotated proximally to the annotated transcript start site, and intronic peaks (Figure 2b). For distal peaks, we observed significant increases in number between early and late stages in the spider B ohnologues, and both species’ single-copy orthologues, and a significant increase between early and middle stages for spider single-copy genes (Figure 2d; Table S1). The number of intronic peaks increased in time, in a similar pattern to distal peaks (Figure 2f; Table S1).

In contrast, TSS peak heights decreased in time, with significant decreases between early and late and/or early and middle stages for harvestman pro-orthologues of ohnologues and both species’ single-copy genes (Figure 2e; Table S1). Mean TSS peak heights were lower for both spider ohnologues later compared to early stages, but this difference was not significant (Figure 2e; Table S1). To confirm this pattern, we investigated whether there might be some genes with no TSS peaks in early or middle stages, meaning they would be excluded from the dataset when the logarithm was taken (i.e., log10 of 0 would return “-Inf”) resulting in them only being counted in late stages and decreasing the median or mean of the late-stage data; however, this did not affect the negative relationship of TSS peak height with time. Any genes without TSS peaks may have misannotated start sites, resulting in no TSS peaks in any stage, rather than no TSS peak in any specific stage.

### Differences between species as well as ohnologue and single-copy orthologue gene families

While differences in ATAC peaks and expression between the stages were small and rarely significant, there were clear differences in these metrics between the species and between gene types when stages were combined. Under our null hypothesis, consistent with the outcome of subfunctionalisation, we predicted that both spider ohnologues would exhibit a decrease in expression to the same or a similar extent, and that the sum of counts for spider A and B ohnologues would be nearly the same as their single copy harvestman pro-orthologues. Consequently, we would also predict that ohnologue expression would be lower than the spider single-copy genes, which we would predict to be equivalent to their harvestman orthologues (and to the harvestman pro-orthologues of the ohnologues). However, we found that the mean of normalised counts for ohnologue A, which we defined as the higher expressed ohnologue of the pair, was 7% lower than that of the harvestman pro-orthologue, while ohnologue B, the lower expressed ohnologue, was 43% lower, so while both ohnologues were lower in expression compared to the harvestman, ohnologue B was usually much lower (Figure 3a; Figure S2; Table S1). We observed that expression was not significantly different between the species’ single-copy orthologues, as expected (Figure 3a; Figure S2).

**Figure 3:**
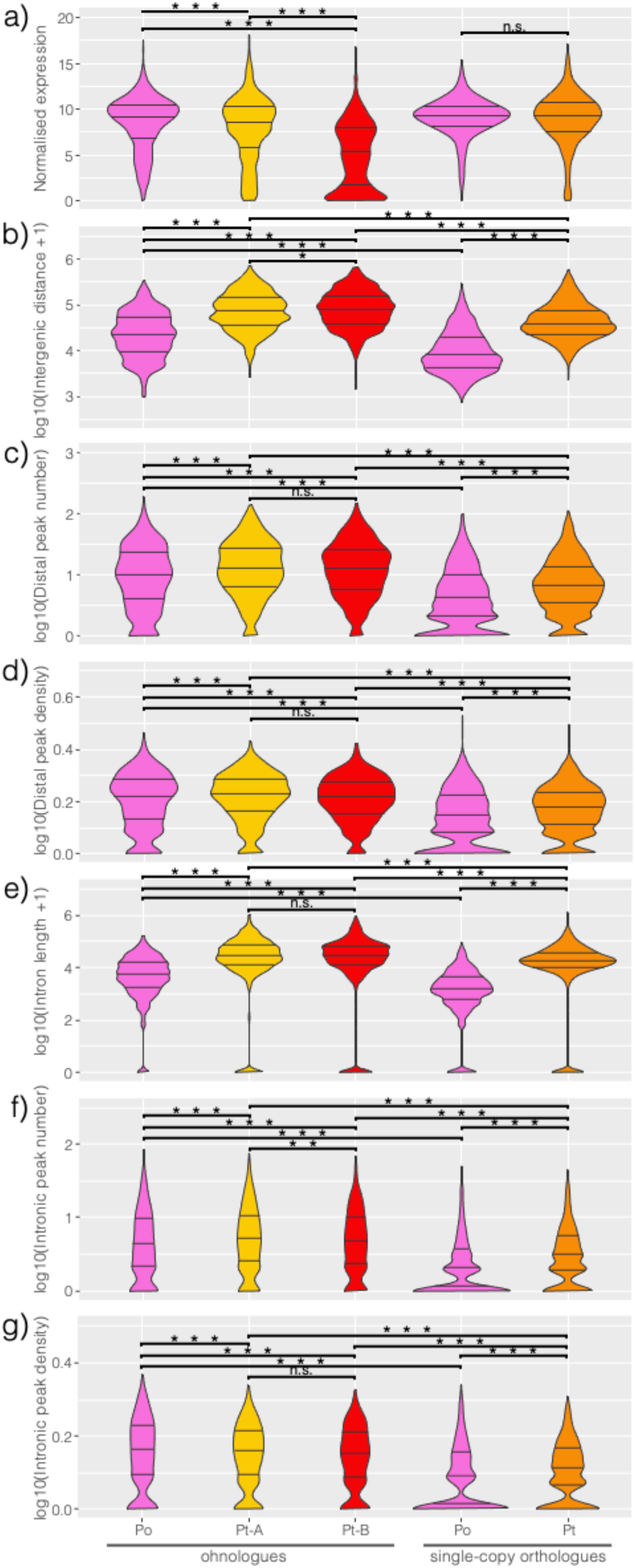
Differences in expression, ATAC peaks, intergenic distances and intron lengths between harvestman and spider genes, and between ohnologues and single-copy orthologues. a) – g) Violin plots of normalised expression counts, intergenic distance, distal peak number and density,intron lengths, and intronic peak number and density per gene, for ohnologue and single copy orthologue families. Significance levels were determined with a two-tailed Wilcoxon rank sum test (*** p<0.001; ** p<0.01; * p<0.05; n.s. p>0.05).

We also noticed differences in gene structure and peak density between the two species, as well as between ohnologues and single-copy orthologues (Figure 3). Firstly, spider genes have longer intergenic distances, more distal peaks, and longer introns than harvestman genes, regardless of belonging to ohnologue or single-copy orthologue families (Figure 3b, c, and e; Table S1). This likely represents differences in genome organisation and size. The *P. tepidariorum* genome is larger than the *P. opilio* genome, with effective genome sizes of 1.09x10^9^ and 4.76x10^8^, respectively.

Spider ohnologues have more intronic peaks than harvestman ohnologue pro-orthologues, and spider single-copy genes have more intronic peaks than harvestman single-copy orthologues, though harvestman ohnologue pro-orthologues had more intronic peaks than spider single-copy genes (Figure 3f). This pattern indicates differences between gene types besides species-specific differences.

Genes in ohnologue families had longer intergenic distances and intron lengths compared to single-copy orthologues, a pattern which held true for their pro-orthologues in the harvestman (Figure 3b, e). To investigate this further, we looked at the density of peaks, both in intergenic and intronic regions. Distal peak density was higher in ohnologue families than single-copy gene families (Figure 3d, Table S1), and was similar between spider ohnologues and the harvestman pro-orthologues, reflecting the higher number of distal peaks observed for spider ohnologues (Figure 3c), but accounting for the smaller intergenic distances in the harvestman (Figure 3b). A similar pattern was observed for intronic peaks, when accounting for differences in intron length between species revealed gene-type differences in intronic peak number (Figure 3g). Genes in ohnologue gene families have higher intronic peak density, with no difference between the A vs B ohnologues, and the harvestman ohnologue pro-orthologues having the highest intronic peak density.

Harvestman single-copy orthologues had the lowest intronic peak density (Figure 3g).

### Temporal subfunctionalisation

We then examined the number or stages each single copy gene or ohnologue was expressed in relative to its harvestman orthologue using the relative expression (log2 fold change) between the stages to determine its stage-specificity. Single-copy orthologues were most likely to be expressed consistently across all three stages in both *P. tepidariorum* and *P. opilio* (Figure 4a). For ohnologues, we examined whether the lower expressed ohnologue B, was detected in the same number of stages as the higher expressed ohnologue A, or in fewer stages. We found that ohnologue B was more likely to be one- or two-stage specific, while ohnologue A was more likely to be expressed in all three stages (Figure 4b). This indicates that the lower expression of the B ohnologue is consistent with a temporal restriction in expression, rather than a uniform reduction in expression levels across all stages. This is particularly evident for *Scr* (Figure 4c.i), where *Pt-Scr-A* is expressed consistently across all three stages, similarly to *Po-Scr*, while *Pt-Scr-B* is expressed at similar levels to *Pt-Scr-A* at only the middle and late stages. Thus, the decrease in expression for *Pt-Scr-B* occurs by temporal restriction to middle and late stages. When both A and B ohnologues are single-stage specific in expression, the B ohnologue is expressed at a lower level, e.g., for *mab21* genes (Figure 4c.ii). Comparing spider *Tspf74F* ohnologues to *Po-Tsp74F* shows that while *Pt-Tsp74F-A* has a higher average expression across the stages, *Pt-Tsp74F-B* has the highest expression of either ohnologue, but only in the early stage (Figure 4c.iii), showing that stage specificity may reflect a gain of expression. Overall, this shows that single-copy genes do not change in expression levels between the developmental stages we investigated, while ohnologues have different temporal expression patterns between stages.

**Figure 4:**
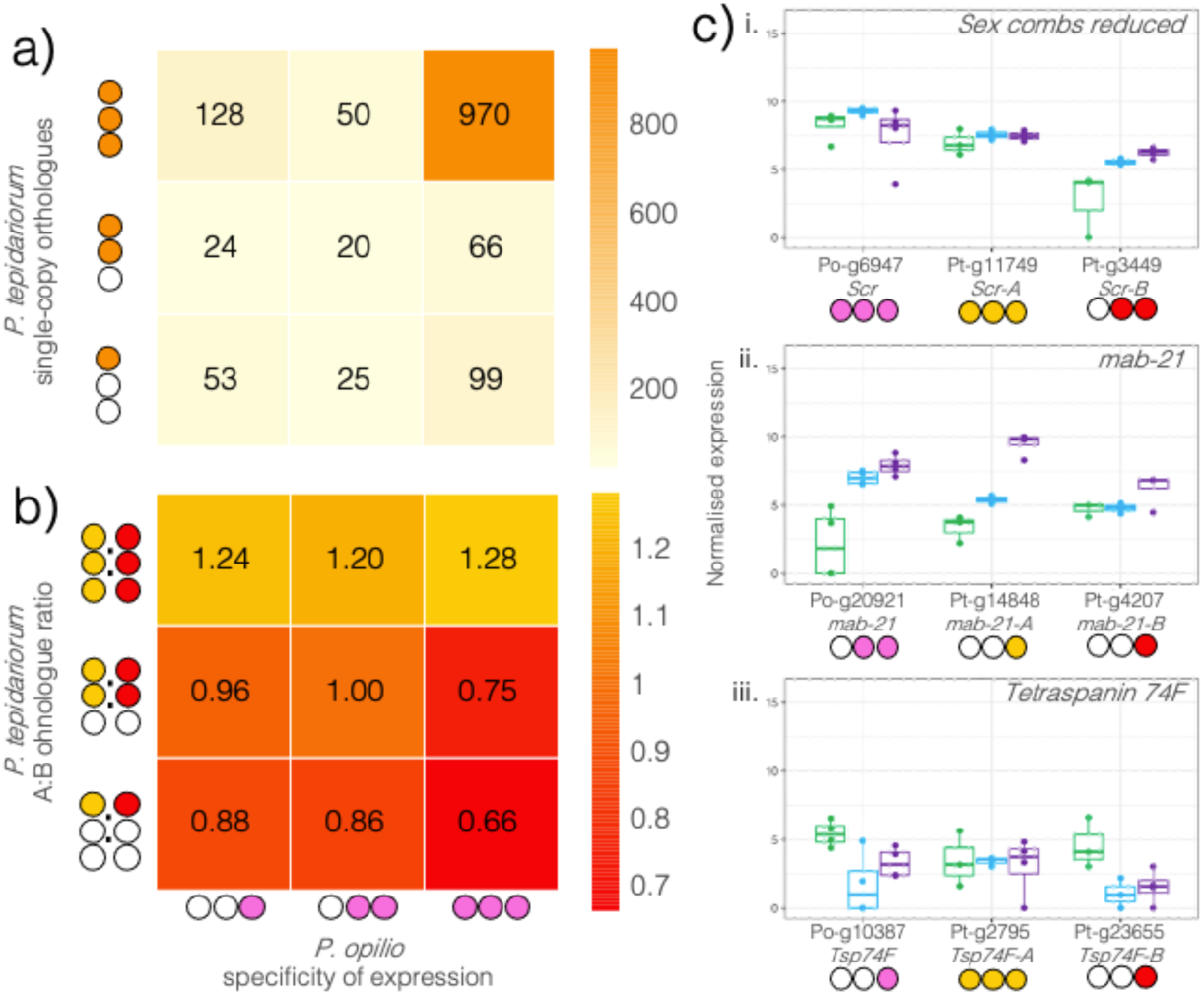
Stage specificity of expression of single-copy orthologues and spider ohnologues. a) Heatmap of the number of spider single-copy orthologues expressed in either a single stage, two, or three stages, by the number of stages their harvestman orthologues are expressed in. b) Heatmap of the ratio of spider A to B ohnologues expressed in either a single stage, two, or three stages, by the number of stages their harvestman pro-orthologues are expressed in. c) Expression of examples of three ohnologue OGs with different stage specificities for the harvestman and spider genes. i. *Scr*, ii. *mab-21*, iii. *Tsp74F*. Circles below the gene IDs indicate the number of stages the gene is expressed in. Gene names reflect previous annotations (Schwager et al. 2017) or annotated *P. tepidariorum* gene names on NCBI.

### Ohnologue and single-copy gene families have different annotated functions

Based on the differences in chromatin accessibility and gene structure between the gene types, we then assessed the function of genes in ohnologue versus single-copy orthologue gene families. GO biological processes for development, morphogenesis, and differentiation were overrepresented among ohnologue families, while housekeeping functions like metabolism, transport, synthesis, and translation were overrepresented among single copy orthologue families (Figure S3a). While single-copy gene families were enriched for the GO biological process of “regulation of transcription by RNA-polymerase II”, the more specific term of “regulation of gene expression” was among the enriched terms for ohnologue families. Similarly, among terms related to protein synthesis, most were associated with single-copy gene families, but proteins annotated with phosphorylation activities were overrepresented among ohnologue families, suggesting ohnologues have functions as both transcription factors and kinases, potentially as components of signalling pathways. We also verified that GO terms were consistent between harvestman and spider reference genome annotations. As expected, GO terms were highly correlated between the species among ohnologue families and among single-copy gene families (Figure S3c.i).

### Early- and late stage-specifically expressed genes

We also identified the genes that were early- or late-specific in their expression patterns, using log2 fold change as we had done for stage specificity. Early-specific genes were overrepresented for biosynthetic processes and intracellular localization, consistent with the functions for early development. Late-specific genes were annotated almost exclusively for GO biological processes and GO molecular functions not found among early-specific genes, including cell adhesion, DNA binding, and Wnt signalling (Figure S3b). We found again that GO annotations were highly correlated within the stage specificity groups and between species, confirming congruence in GO annotation between the two genomes (Figure S3c.ii).

### Spatial expression of late-expressed genes

Given the developmental functions annotated to late-specific genes, we then used single-cell data from partially overlapping developmental stages (stages 7, 8, and 9, compared to our stages 5, 7, and 8) in *P. tepidariorum* (Leite et al. 2024) to determine whether spider genes that were late-specific in expression and had late-specific ATAC peaks were associated with any particular cell clusters. We used differential peak numbers from comparisons between the three stages to determine if a gene was annotated with more peaks in the late-stage relative to the early and middle stages. We found for most of the spider late-specific genes that the harvestman orthologue was also expressed most highly in the late stage, for example, the Toll ligand *Po-spz3* (the harvestman orthologue of fly *spatzle 3*), *Po-HSPA12A* (the harvestman orthologue of *Heat shock 70kDa protein 12A* which seems to lack a fly orthologue (Wada et al. 2006)), *Po-lin-28* (the harvestman orthologue of fly *lin-28*), and *Po-BAMBI* (the harvestman orthologue of *BMP and activin membrane-bound inhibitor* which seems to lack a fly orthologue); exceptions were *Po-Pitx* (the harvestman orthologue of *Pituitary homeobox*, orthologous to fly *Ptx1*) and *Po-FoxP* (the harvestman orthologue of fly *Forkhead box P*) (Figure 5a.i-ix).

**Figure 5:**
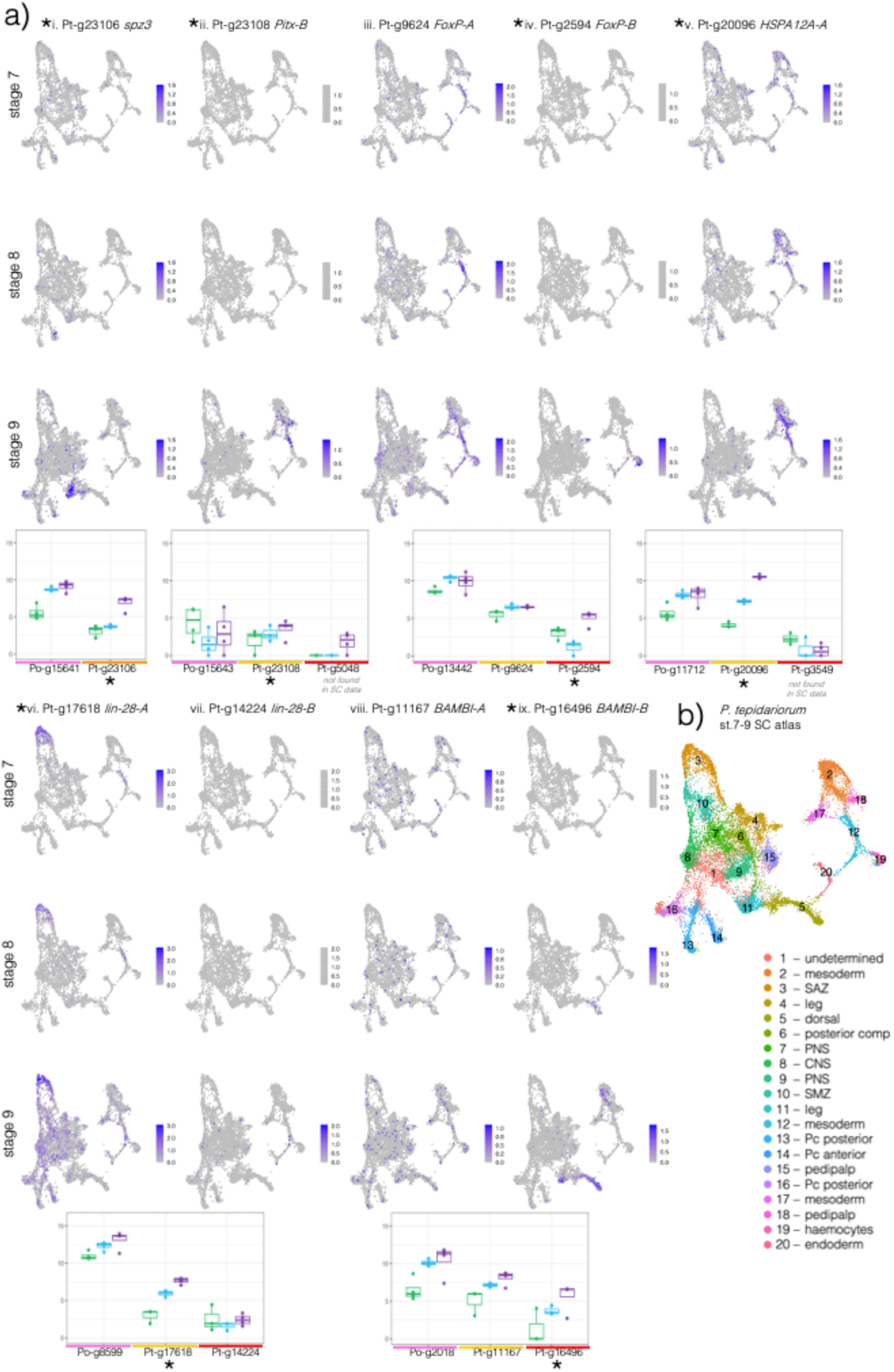
Expression of late-specific spider genes, their ohnologues, and harvestman orthologues in our data and the single-cell atlas from Leite *et al*. (2024). Genes that were late-specific in their expression and had late-specific peaks are marked with asterisks. Gene names are based on the annotated spider name on NCBI or the closest fly orthologue, except for *Pitx-A* and *Pitx-B* which were previously identified in Aase-Remedios *et al*. (2023) and *FoxP-A* and *FoxP-B* which were previously identified in Janssen *et al*. (2022).

Of the 25 genes we identified with both late-specific expression and late-specific peaks, 20 belonged to ohnologue families. Some of these ohnologues were found to be present in specific cell clusters (Figure 5b), namely *Pt-Pitx-B*, *Pt-FoxP-B*, *Pt-HSPA12A-A*, *Pt-lin-28-A* and *Pt-BAMBI-B* (Figure 5.ii-vi, and ix), though the rest were widely expressed and simply increased in expression levels over time (Figure S4). A few had ohnologues that were not present in the single cell data, e.g., *Pt-Pitx-A* and *Pt-HSPA12A-B*, but for the others we could compare the expression of the ohnologues to detect spatial as well as temporal subfunctionalisation.

Harvestman *FoxP* and spider *FoxP-A* are expressed at similar levels in middle and late stages, and only *FoxP-B* is late-specific (Figure 5a.iii-iv). Both spider ohnologues are expressed at lower levels relative to harvestman *FoxP*, consistent with subfunctionalisation. *FoxP* ohnologues are also found in different cell clusters, with *FoxP-A* expressed in cells annotated as the mesoderm (cluster 12) and central nervous system (CNS, cluster 8) (Figure 5a.iii) and *FoxP-B* in cells annotated as haemocytes (cluster 19) and some in the developing legs (cluster 4) (Figure 5a.iv). This is largely consistent with published expression data where *FoxP-A* was detected in later stages (10+) in the CNS, with stage 8 to 9 head expression that could represent mesodermal cells in cluster 12, and *FoxP-B* was detected in haemocytes (*FoxP-B* is *FoxP1*; *FoxP-A* is *FoxP2*) (Janssen et al. 2022).

The differences between *Po-Pitx* expression and *Pt-Pitx-A* and *Pt-Pitx-B* appears to reflect both the subfunctionalisation of the spider ohnologues and differences in the timing and expression patterns between the species, with *Po-Pitx* expressed highest in the early stage, lowest in the middle, and then slightly higher in the late stage, while both spider ohnologues are highest during the late stage (Figure 5a.ii-iii). *Pt-Pitx-B* was detected in cells annotated as the mesoderm (clusters 12 and 2) and the pedipalps (cluster 18) (Figure 5a.ii). *Pt-Pitx-A* was not detected in early or middle stage embryos, indicating it may be activated later than *Pitx-B*, and it was not detected in the single-cell data, suggesting it is only very lowly expressed until stage 9. This is largely consistent with expression data from later stages (stages 10 to 12) showing that *Po-Pitx* and *Pt-Pitx-B* are both expressed in the pre-cheliceral region, while similar patterns to the expression of *Po-Pitx* along the ventral midline are partitioned between *Pt-Pitx-A* and *Pt-Pitx-B* (Leite et al. 2018). Note that we are using names as in Aase-Remedios et al. (2023), so, in this case, *Pitx-A* is the lower-expressed ohnologue; *Pitx-A* is *Pitx-1*, *Pitx-B* is *Pitx-2* in Leite et al. (2018).

Expression differences consistent with subfunctionalisation were also seen for the other late-specific genes with retained ohnologues. *P. tepidariorum lin-28-A* is expressed in cells annotated as the segment addition zone (SAZ) at stage 7, 8 and 9, though expression was also detected across many other cell clusters at stage 9 (Figure 5a.vi). *Pt-lin-28-A* is also expressed at a low level in the cells annotated as cluster 12 (mesoderm) throughout all these stages. Its ohnologue *Pt-lin-28-B*, is not expressed at stages 7 and 8, and expression was at a much lower level at stage 9, with its highest expression in cluster 12, and it was detected in a few cells that also express *lin-28-A* (Figure 5a.vii). *Pt-lin-28-B* may be expressed in later stages relative to *Pt-lin-28-A*, but it is clear that *lin-28-B* is expressed in fewer cells and primarily in different cell clusters relative to *Pt-lin-28-A*.

### Double conserved synteny of a late-specific locus

We noticed that two of the late-specific genes, *Pt-Pitx-B* and *Pt-spz3* were located one gene apart, and so were the harvestman orthologues of these genes (*Po-Pitx* and *Po-spz3*), indicating there are two conserved regions consisting of three genes in each species. This prompted us to identify the late-specific peaks in each *spz3 – Pitx* locus and investigate the synteny surrounding these loci (Figure 6).

**Figure 6:**
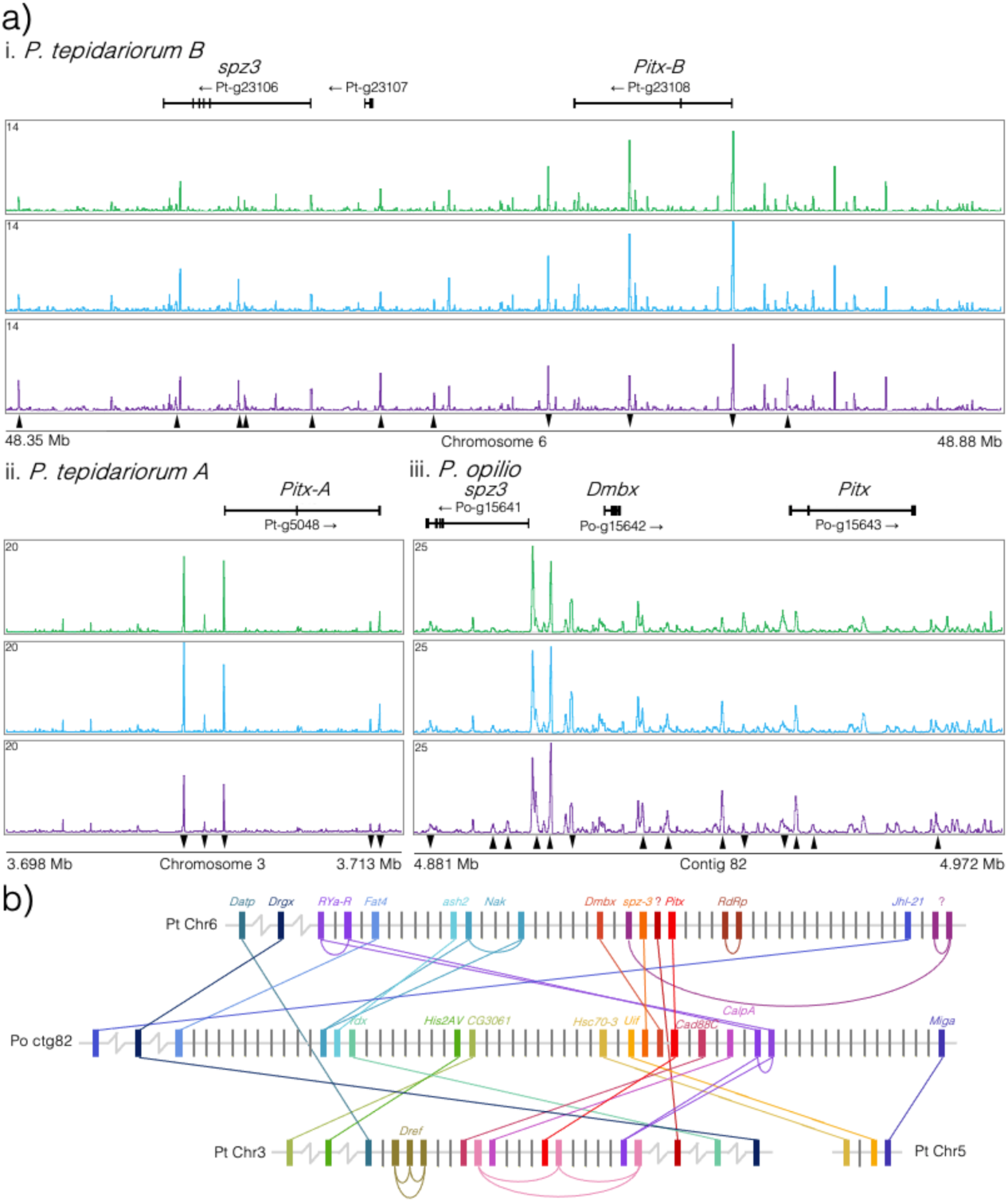
Chromatin profiles and synteny surrounding the conserved late-specific *spz3* – *Pitx* locus in *P. tepidariorum* and *P. opilio*. a) Chromatin accessibility profiles for i. the *Pt-spz3 – Pt-Pitx-B* locus on chromosome 6 in the spider, ii. The *Pt-Pitx-A* locus on chromosome 3 in the spider, and iii. the *Po-spz3 – Po-Pitx* locus on contig 82 in the harvestman. Black arrowheads indicate the significant DiffBind change in peak height for peaks that are higher in the late stage (pointing up) or early stage (pointing down). b) Schematic of gene families with members linked to *Pitx* loci in the two species. Families are coloured according to their relative position along the harvestman contig 82. Names represent previous annotations (Aase-Remedios et al. 2023) or the name of the fly orthologue and question marks indicate uncharacterised genes without an identified fly orthologue.

We identified several late-specific peaks near the promoter of *spz3* and one upstream of *Pitx-B*, which correlate with the late-specific expression of these genes in the spider (Figure s7a.i and 6a.ii). Within the second intron of *Pitx-B* we also identified a peak that was high at both early and middle stages but lower at the late stage, in a pattern directly inverse to *Pitx-B* expression (Figures 7a.i and 6a.ii). In the ohnologous *Pitx-A* locus, we did not identify any late-specific peaks, rather all peaks surrounding this gene were lower in the late stage relative to the early or middle stages (Figure 6a.ii). In the harvestman, we identified a peak proximal to the promoter of *Po-Pitx* that was highest in the early stage, lowest in the middle stage, and increased again from middle to late, mirroring the expression pattern observed for *Po-Pitx* in our data (Figures 7a.iii and 6a.ii).

**Figure 7:**
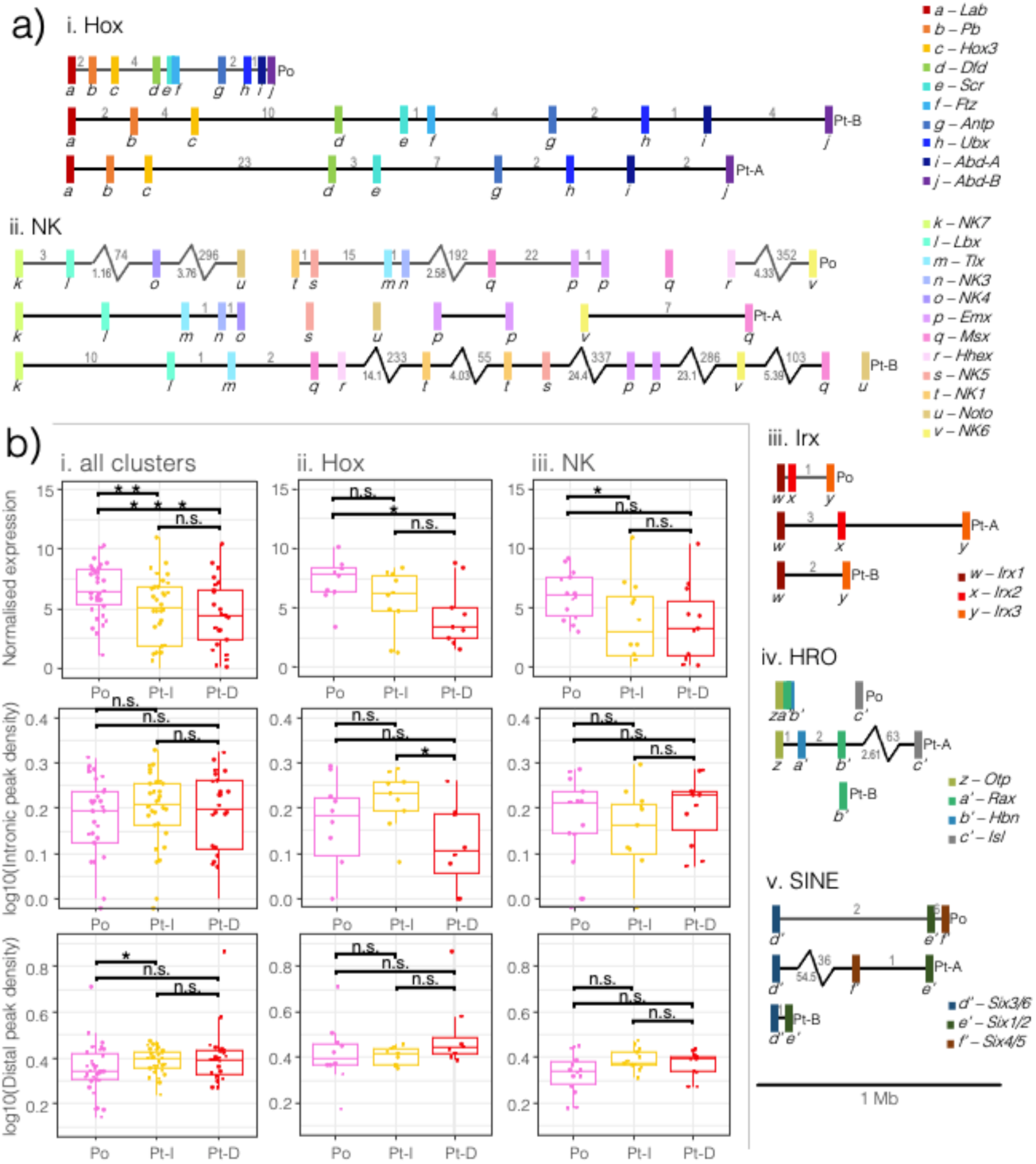
Homeobox gene organisation, expression, and ATAC peak density metrics in the spider and harvestman. a) To-scale schematic of conserved homeobox gene clusters in the spider and harvestman: i. Hox, ii. NK, iii. Irx, iv. HRO, and v. SINE. Black or grey lines represent chromosomes and contigs, respectively. Coloured rectangles indicate homeobox genes, plotted by the position of the homeobox; letters below and the colour scheme indicate the gene family, with the key to the right. Numbers above the chromosome/contig lines indicate the number of genes between each homeobox gene on either side. The zig-zag bar indicates a long distance, the length of which in Mb is denoted below in grey. The full annotation of *P. opilio* homeobox genes can be found in Supplementary file S2. b) Expression, intronic peak density, and distal peak density for i. all homeobox clusters depicted in a), and for ii. just Hox and iii. Just NK genes. Pt-I refers to the more intact cluster (cluster A for all clusters besides Hox, as Hox-B is the more intact cluster, after Schwager et al. (2017)), and Pt-D refers to the more dispersed cluster (cluster B for all clusters besides the Hox).

The spider *Pt-spz3 – Pt-Pitx-B* locus is nearly six times larger than the *Po-spz3 – Po-Pitx* locus in the harvestman, comprising 0.53 Mb compared to 0.09 Mb, and *Pt-Pitx-B* is inverted relative to *Po-Pitx* (Figure 6a.i and iii). The intervening gene in the harvestman is another homeobox gene, *Po-Dmbx* (*Diencephalon/ mesencephalon homeobox*, no fly orthologue) (Figure 6a.iii), the spider orthologue of which can be found five genes away (∼0.36 Mb) from *Pt-Pitx-B* on chromosome 6 (Figure 6b). We searched for other homeobox gene families with a similar linkage pattern and found that *Drgx* (*Dorsal root ganglion homeobox*, orthologous to fly *Drgx*) genes are linked to *Po-Pitx* in the harvestman, as well as *Pt-Pitx-B* on chromosome 6 and *Pt-Pitx-A* on chromosome 3 in the spider.

We then searched for genes within 20 genes on either side of each *spz3 – Pitx* locus in harvestman or spider that have orthologues or ohnologues linked to any of the other two *spz-3 – Pitx* loci. The *Pt-Pitx-A* locus does not contain an ohnologue of *spz3*, but there are 12 gene families connecting any two of the three *spz3 – Pitx* loci (the three loci comprising the contig bearing the harvestman *spz3 – Pitx* locus and the two spider chromosomes containing *spz3 – Pitx-B* or *Pitx-A*), while 3 families connect the harvestman *Pitx* locus to spider chromosome 5, indicating an inter-chromosomal rearrangement (Figure 6b). All the arachnids surveyed in Aase-Remedios et al. (2023) also exhibit linkage between *Pitx*, *Drgx*, and *Dmbx* genes (Supplementary File S1) on two or more chromosomes in the spiders, indicating these genes were linked in the arachnid ancestor, and this linkage was conserved in two ohnologous loci following the WGD.

### Homeobox gene cluster organisation in the harvestman P. opilio

We also examined the chromatin accessibility surrounding homeobox genes in conserved clusters and sub-clusters. We previously characterised the homeobox gene complement of many arachnids, and we found that ancient homeobox gene clusters like the Hox, NK, SINE, HRO, and Iroquois (Irx) clusters were retained with one more intact copy, while the second copy underwent varying degrees of dispersal via gene losses and rearrangements following the arachnopulmonate WGD (Aase-Remedios et al. 2023). Here, we classified the homeobox genes in our improved reference genome assembly for *P. opilio* with the aim of comparing the organisation of homeobox gene clusters to the expression and chromatin accessibility between clusters with differing levels of organisation (Figure 7a). We hypothesised that intact homeobox clusters might have smaller intergenic distances, more peaks, or differences in expression compared to the more dispersed clusters. Expression is significantly lower for spider homeobox genes than for harvestman genes, but spider homeobox genes have higher intronic and distal peak densities (Figure 7b.i).

We then compared only the Hox and NK clusters, two ANTP-class clusters with different levels of conservation and organisation (Aase-Remedios et al., 2023). The harvestman Hox cluster is compact and organised, while both spider Hox clusters are organised, but larger and have more intervening genes, and even more so for the Hox-A cluster, which has lost *ftz*, making it the more dispersed of the two (for all other clusters, the B cluster is more dispersed) (Figure 7a.i). The NK genes are organised to a similar extent in the harvestman and spider, comprising smaller sub-clusters of two or three NK genes separated by large distances, onto different contigs, or even chromosomes (Figure 7c.ii). The harvestman NK cluster is found on four contigs, one with *NK7*, *Lbx*, *NK4*, and *Noto*, another with *NK1*, *NK5*, *Tlx*, *NK3*, *Msx1*, *Emx1*, and *Emx2*, a third with *Msx2*, and a fourth with *Hhex* and *NK6* (Figure 7c.i). Some clustered gene pairs in the harvestman mirror the conserved clustering in the spider, indicating that sub-clustering of certain NK gene pairs is constrained in both species. These conserved pairs consist of *Po-NK7 – Po-Lbx* and *Pt-NK7-A – Pt-Lbx-A*, *Po-Tlx – Po-NK3* and *Pt-Tlx-A – Pt-NK3*, *Po-NK1 – Po-NK5* and *Pt-NK1-2 – Pt-NK5-B*, and *Po-Emx1 – Po-Emx2* and both *Pt-Emx1-A – Pt-Emx2-A* and *Pt-Emx1-B – Pt-Emx2-B*.

Hox-A cluster expression was significantly lower than that of the genes in the harvestman Hox cluster, and lower than the expression of intact cluster genes in Hox-B, though this decrease was not significant (Figure 7b.ii). This pattern mirrors the difference in expression between harvestman pro-orthologues and spider ohnologues (Figure 3a). For the NK cluster, the intact copy which contains the more conserved ‘core’ NK cluster (NK-A) was significantly lower-expressed than the harvestman NK genes, while the more dispersed NK-B was also lower, but not significantly different from the harvestman (Figure 7b.iii). The intact Hox-B cluster also had significantly higher intronic peak density than the dispersed Hox-A cluster and was not significantly higher than the harvestman Hox cluster intronic peak density. (Figure 7b.ii). There were no significant differences in peak density metrics between the NK clusters.

The other homeobox gene clusters do not contain enough genes to reveal any discernible patterns in expression or peak metrics. However, they have varying degrees of organisation relative to the spider clusters (Figure 7c). The harvestman Irx cluster is smaller than either of the spider clusters, and the HRO cluster of the harvestman is extremely compact relative to the spider. Both the harvestman HRO and SINE clusters are inverted relative to the spider clusters, though in which lineage this occurred cannot be determined. The spacing of *Six3/6* further away from *Six1/2* and *Six4/5* in the harvestman matches that seen in the intact *P. tepidariorum* SINE-A cluster, and the harvestman Irx cluster is similarly proportionally spaced to the *P. tepidariorum* Irx-A cluster as well. These organisations illustrate that the diversity of levels of organisation observed between spider homeobox gene clusters are also present in the smaller *P. opilio* genome. To-scale figures of conserved core clusters and sub-clusters including intervening genes, with chromatin profiles can be found in Supplementary file S3.

## Discussion

The data we present here adds new depth to our understanding of the impact of the arachnopulmonate WGD on gene expression and regulation during development. We found that spider ohnologues are expressed at lower levels relative to their harvestman pro-orthologues, with one ohnologue typically being lower expressed than the other, indicating an asymmetrical result of subfunctionalisation. Under the standard model of subfunctionalisation, we expected a decrease in the number of peaks in the lower-expressed ohnologue consistent with the loss of regulatory elements (Force et al. 1999), while under specialisation, we would expect an increase consistent with the fine-tuning of regulation (Marlétaz et al. 2018). We did not find any significant differences in the regulatory complexity (i.e., peak number or peak density) between ohnologues in the spider, despite an asymmetrical decrease in expression between ohnologues (Figure 3).

We also observed different dynamics for TSS peaks compared to distal and intronic peaks over time (Figure 2). TSS peak heights decreased over time, and this difference was significant for all genes except spider ohnologues. Distal peaks and intronic peaks, however, increased significantly in number in the spider, and showed a non-significant increase in the harvestman. Distal peaks are likely more dynamic, representing potential enhancers and repressors, thus the increases in distal peak numbers may represent the gradual opening of chromatin at *cis*-regulatory elements as development proceeds. The decrease in TSS peak heights amid the opening of more distal and intronic peaks may reflect that as development proceeds, more of the genome is in regions of open chromatin, causing a dilution of the Tn5 transposase relative to the amount of accessible genome sequence, resulting in shorter constitutive peaks (like those at promoters) due to the increase in new peaks.

There were interesting differences between the two species and between ohnologues versus single-copy genes in how chromatin dynamics changed with time, suggesting these genes might be structurally different. When we examined intron size, intergenic distances, and intronic and distal peak number and density for combined stages, we found differences in regulatory complexity and expression between the gene families with retained ohnologues relative to those that returned to the single copy. While the spider has a larger genome, so we would predict longer introns and intergenic distances, we also saw that spider and harvestman genes in ohnologue gene families had longer introns and intergenic distances than those in single copy gene families, but also had higher distal and intronic peak density than either species’ genes in single-copy gene families. Therefore, the arachnopulmonate WGD seems to have impacted the genome structure of the spider beyond duplicate gene retention, resulting in longer introns and larger intergenic distances, which is also true of vertebrate genomes relative to outgroups, where gene number does not increase proportionally with genome size (Ohno 1999). Underlying the WGD, we can infer that the genes that were later retained in duplicate in the spider after WGD had higher regulatory complexity in the spider-harvestman ancestor, because ohnologue pro-orthologues in the harvestman have higher peak density than the harvestman single-copy orthologues.

Ohnologues were enriched for developmental functions, including regulation of transcription and signalling pathways, consistent with previous findings (Putnam et al. 2008; Berthelot et al. 2014). Our conservative definition of ohnologues to enable our two-species comparisons here identified 530 ohnologous gene families between *P. tepidariorum* and *P. opilio*, comprising 530 harvestman genes and 1,060 spider genes. This is likely an underestimation of the true number of spider ohnologues originating from the arachnopulmonate WGD and should not be interpreted as a count of all the ohnologues in *P. tepidariorum*. Relaxing our definitions to allow for independent tandem duplications or gene losses in the outgroup or the three spiders we used to define them, as well as rearrangements bringing ohnologues onto the same chromosome, would increase the number of detected ohnologue gene families. We also compared early- and late-specifically expressed genes and found that late-specific genes were enriched for interesting developmental functional annotations.

We examined genes that were expressed specifically in the late-stage RNA data and had corresponding distal peaks that increased in the late-stage ATAC data, as these would be genes expressed during the formation of the germ band and the onset of segmentation. 20/25 of these late-specific genes belonged to families that had retained ohnologues and included several interesting developmental genes (Figure 5 and S4). Two of the late-specific genes were found in a conserved arrangement in both the spider and harvestman genomes and revealed a pattern of double-conserved synteny resulting from the WGD (Figure 6). In the harvestman *Pitx* was not late-specific in expression, but *Po-spz3* was (Figure 5a.i-ii).

Gene linkage is thought to be constrained by shared regulatory elements (Irimia et al. 2012), but here, it appears that there has been an inversion in one of the lineages, which may have affected any co-regulation between these genes. This could relate to how in the spider, *Pt-Pitx-B* and *Pt-spz-3* are in the same orientation in the genome and are both late-specific in expression, while the two genes are in the opposite orientation to one another in the harvestman and have different temporal expression patterns, though this remains to be tested.

What was also apparent from the conservation of this locus was a pattern of 1:2 synteny resultant from the arachnopulmonate WGD. Many genes linked to the harvestman *spz3* and *Pitx* genes could be found on chromosome 6 or chromosome 3 in the spider, which bear the two *Pitx* ohnologues. While there are many gene families with only 1:1 relationships among harvestman and spider *Pitx* loci, this pattern still illustrates the 1:2 conserved synteny resulting from the WGD, as single-copy orthologues are found linked to one or the other of the two *Pitx* ohnologues. WGD syntenic patterns can be obscured by gene losses or rearrangements. Our synteny analysis included gene families that underwent the loss of a harvestman pro-orthologue (e.g., the paralogs *Pt-g23017* and *Pt-g5092* (uncharacterised family, adjacent to *Pt-Pitx-B* and linked to *Pt-Pitx-A*, respectively)) and a spider ohnologue (e.g., *Pt-spz3* and *Po-spz3*, as well as the rearrangement onto another chromosome, in this case (e.g., *Po-Uif* and *Pt-Uif* on chromosome 6) (Figure 5b). This is consistent with the patchwork arrangement of vertebrate ohnologues across four paralogous chromosomes due to gene losses and rearrangements (McLysaght et al. 2002; Dehal and Boore 2005; Singh et al. 2015). This is also consistent with findings that the 1:2 pattern of synteny is present, but somewhat obscured in spider genomes due to extensive rearrangements, but are apparent among homeobox gene clusters (Fan et al. 2021; Aase-Remedios et al. 2023; Miller et al. 2023; Kulkarni et al. 2024).

With our improved genome assembly, we have now described the homeobox gene complement and organisation in the harvestman (Figure 7a and Supplementary File S2), and enabled better detection of the linkage between homeobox genes compared to previous assemblies (Gainett et al. 2021). We confirmed that there is a single intact Hox cluster containing all 10 genes and found that the other homeobox clusters exhibit different levels of organisation in *P. opilio*, with a very condensed HRO cluster and similar spacing of genes in the SINE and Irx clusters in spiders and the harvestman. The harvestman NK genes can be found in dispersed sub-clusters, some of which are conserved in two copies in the spider, though without chromosome-scale resolution, it is not clear if some of the NK sub-clusters are linked. We previously described the expression of NK cluster genes in the harvestman and found that harvestman *NK7*, *Lbx*, *Tlx* and *NK3* are all expressed in the limbs, and the two *Emx* paralogues are expressed in nearly-identical patterns (Leite et al. 2018; Aase-Remedios et al. 2023), which may implicate *cis*-regulatory elements in the co-regulation and constraint on the organisation of these sub-clustered genes. We were unable to identify any conserved non-coding sequence associated with any peaks in NK sub-clusters between the species due to their evolutionary divergence, so this remains to be investigated. The similarities between the organisation of homeobox genes and the conserved synteny of the late-specific *spz3 – Pitx* locus indicates that the genome of the harvestman may be an appropriate single-copy outgroup for future chromosome-scale syntenic comparisons, more so than other non-arachnopulmonate outgroups like ticks or mites. However, further genome-wide comparisons of macrosynteny relative to ancestral linkage groups are necessary to determine the extent of syntenic conservation in the harvestman relative to other arachnids.

## Conclusions

Our analysis of regulatory regions and gene expression between equivalent embryonic stages of a spider and a harvestman provides evidence that the WGD has impacted the regulation of duplicated genes, many of which have roles in development. Specifically, we showed that ohnologues have likely undergone subfunctionalisation, were characterised by higher regulatory complexity relative to single-copy genes and were enriched for GO functions consistent with roles in development and signalling. This may reflect the propensity of developmental genes, which are thought to have more complex *cis*-regulation, to be retained in duplicate, and is consistent with observations in vertebrates. Using our new reference genome for the harvestman *P. opilio*, we were able to examine the synteny of homeobox genes and a late-specific locus, further providing evidence for the arachnopulmonate WGD. This resource improves upon available genomes from pre-duplicate arachnids to better understand the ancestral state giving rise to both Opiliones and arachnopulmonates, and the impact of the arachnopulmonate WGD.

## Acknowledgements

This work was funded in part by a Leverhulme Trust Grant (RPG RPG-2022-018) to A.P.M. and a Durham-Uppsala Strategic Development Fund award (1875810) to A.P.M. and R.J. We would like to thank Ignacio Maeso for discussion of the analysis.

## Data availability

The sequence data underlying this article are available in GenBank and can be accessed with the accession numbers cited in the Methods upon publication. All code underlying this article is available on GitHub [https://github.com/madeleineaaseremedios/popt_code] and will be made public upon publication.

## Notes

### Competing Interest Statement

The authors have declared no competing interest.

