## Supplementary Figures and Tables for "Assessing the impact of whole genome duplication on gene expression and regulation during arachnid development"

### Supplementary Tables and Figures

Table S1: Means and standard errors of expression and chromatin accessibility metrics for ohnologue and single-copy orthologue gene families by stage, and with stages combined, as well as intron length and intergenic distance by gene type.

| | stage | normalised counts | | TSS peak height | | distal peak number | | intron length | intergenic distance |
|---|---|---|---|---|---|---|---|---|---|
| Po | E | 8.51±0.131 | 8.46±0.0736 | 5.88±0.122 | 5.24±0.0678 | 15.1±0.827 | 16.0±0.503 | 3.61±0.0229 | 4.35±0.0123 |
| | M | 8.53±0.136 | | 4.87±0.100 | | 15.9±0.851 | | | |
| | L | 8.34±0.115 | | 4.97±0.119 | | 17.1±0.930 | | | |
| Pt-A | E | 7.72±0.151 | 7.85±0.0896 | 6.97±0.166 | 6.92±0.0962 | 17.9±0.784 | 19.6±0.507 | 4.20±0.0323 | 4.86±0.0104 |
| | M | 7.91±0.159 | | 7.27±0.173 | | 19.7±0.878 | | | |
| | L | 7.93±0.156 | | 6.54±0.159 | | 21.1±0.956 | | | |
| Pt-B | E | 4.72±0.151 | 4.80±0.0905 | 6.94±0.190 | 6.88±0.106 | 17.0±0.761 | 18.9±0.493 | 4.12±0.0340 | 4.89±0.0103 |
| | M | 4.79±0.164 | | 7.07±0.179 | | 19.2±0.864 | | | |
| | L | 4.88±0.156 | | 6.63±0.180 | | 20.5±0.922 | | | |
| Po | E | 9.09±0.0576 | 9.04±0.0321 | 6.34±0.0706 | 5.69±0.0383 | 7.23±0.311 | 7.52±0.187 | 3.02±0.0159 | 3.97±0.00706 |
| | M | 9.02±0.0576 | | 5.21±0.0551 | | 7.58±0.324 | | | |
| | L | 9.00±0.0515 | | 5.47±0.0666 | | 7.74±0.336 | | | |
| Pt | E | 8.90±0.0753 | 8.93±0.0430 | 6.68±0.0897 | 6.70±0.0529 | 9.85±0.322 | 10.7±0.205 | 3.97±0.0191 | 4.63±0.00576 |
| | M | 8.97±0.0733 | | 6.96±0.0902 | | 10.8±0.353 | | | |
| | L | 8.92±0.0751 | | 6.45±0.0934 | | 11.5±0.383 | | | |

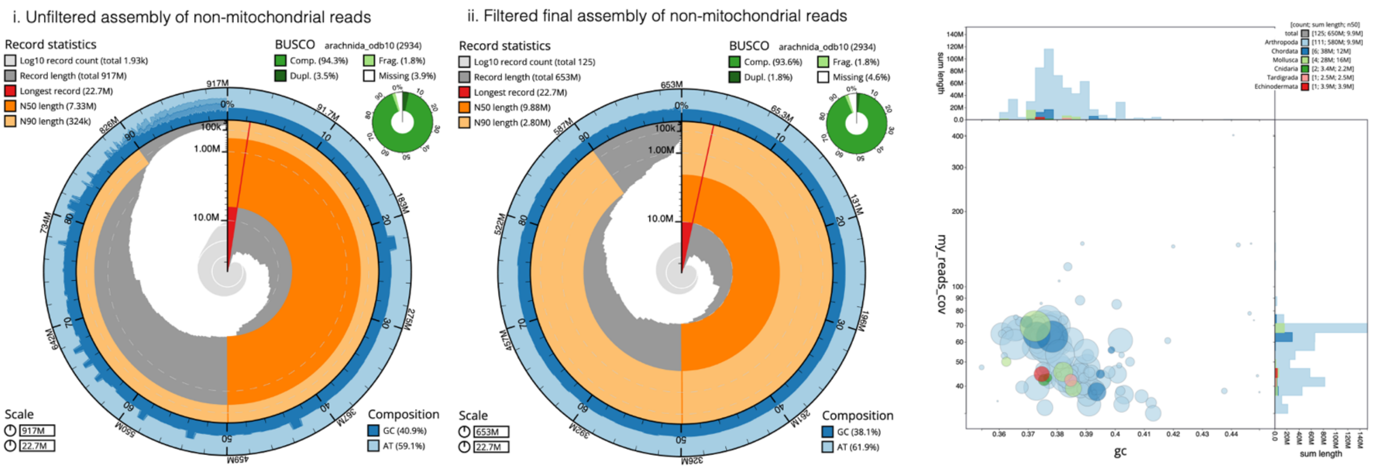


Figure S1: Comparison of the unfiltered and the final filtered assemblies for *P. opilio*. Plots are snail plots made with BlobTools2. Right shows the blobplot of the filtered assembly, plotting the read coverage of each contig by GC content, coloured by phylum of blast hits.

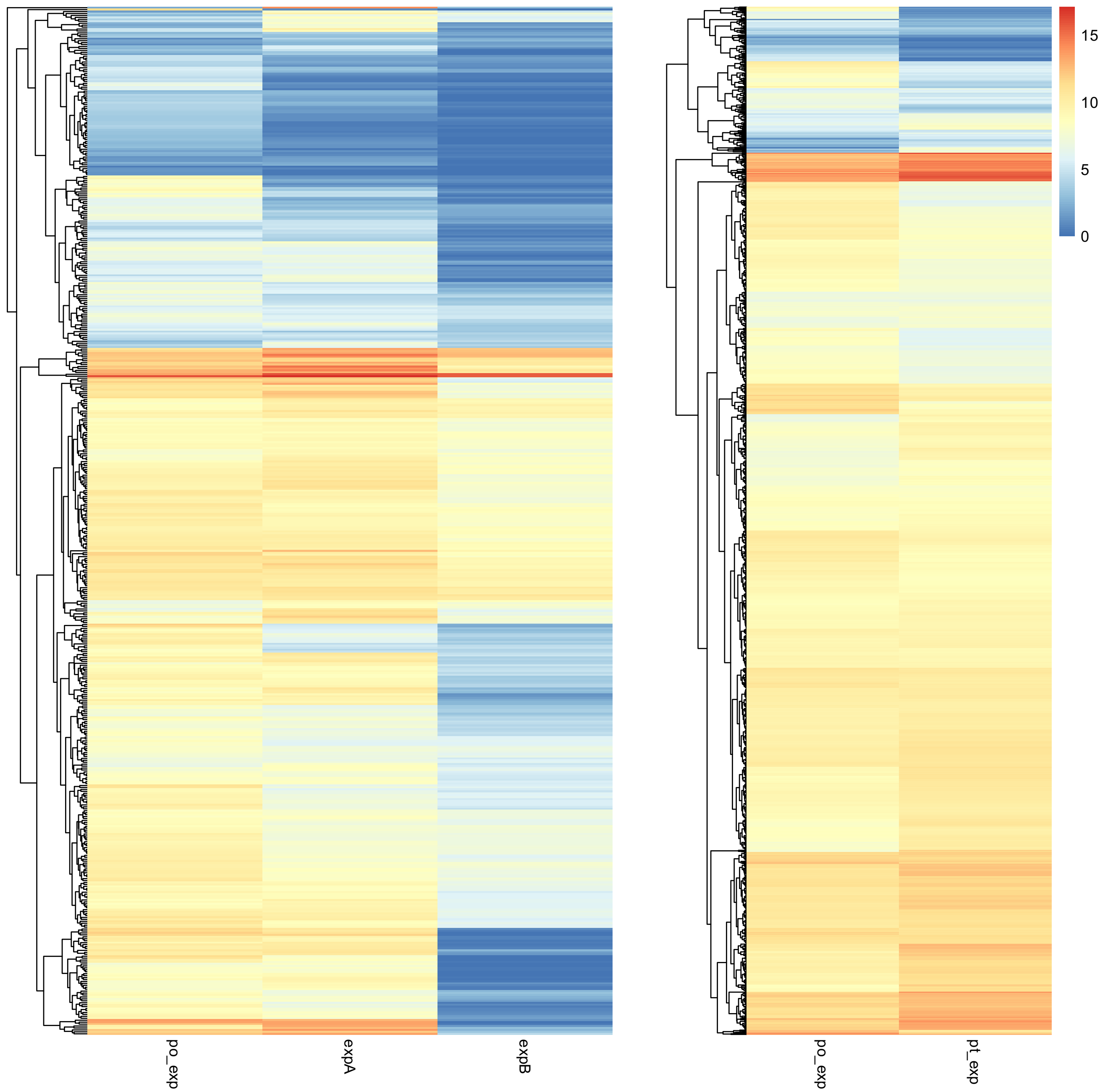


Figure S2: Heatmaps of normalised expression of a) genes in ohnologue families, and b) genes in single-copy gene families. Left is the expression of the harvestman genes (po_exp), right is the expression of spider ohnologues (expA and expB) or orthologues (pt_exp).

9

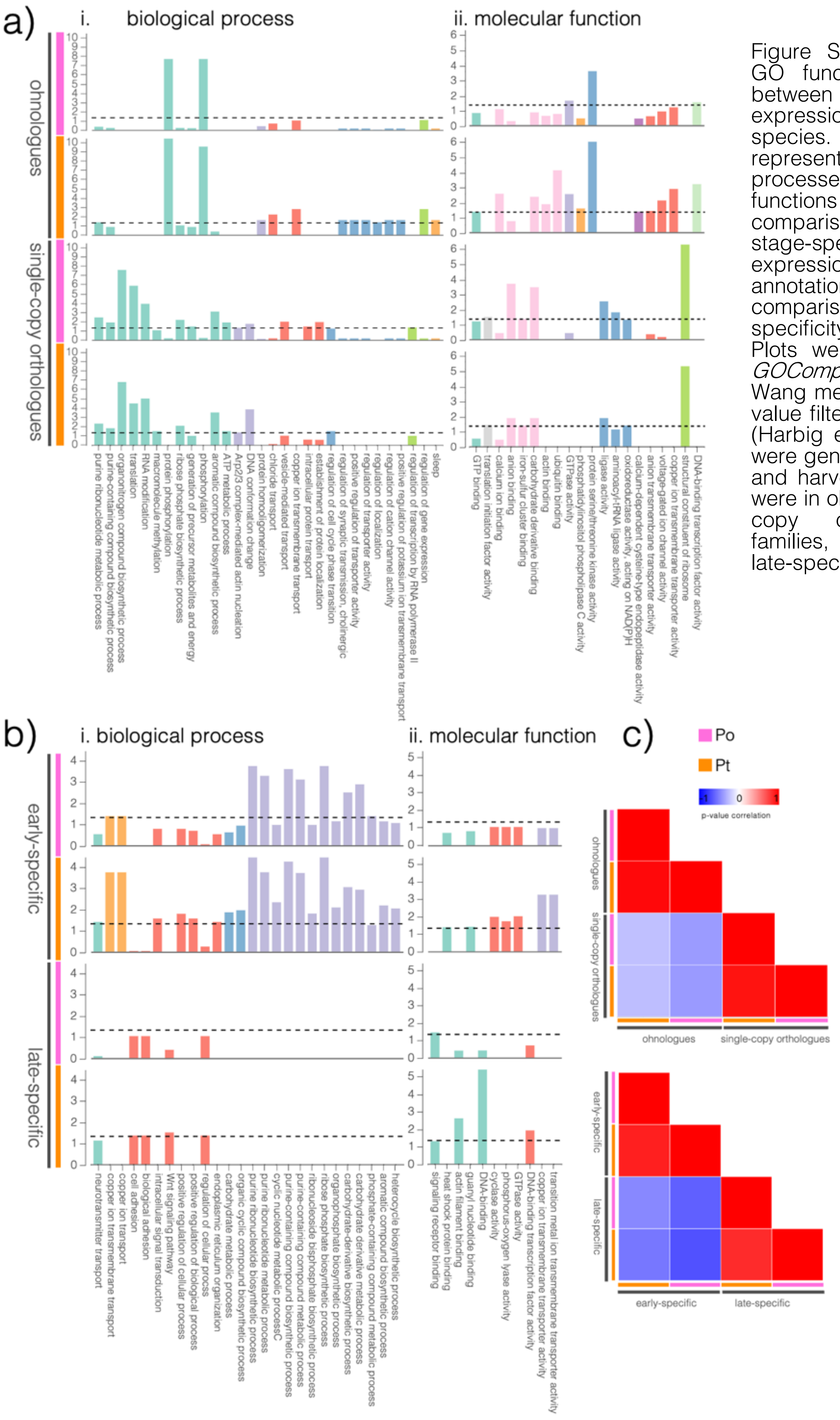


Figure S3: Comparison of GO functional annotations between gene types, expression patterns, and species. a–b) Over-represented GO biological processes (i.) and molecular functions (ii.) for the two comparisons (gene type and stage-specificity of expression). c) Correlation of annotations for (i.) gene-type comparison and (ii.) stage-specificity of expression. Plots were generated with *GOCompass* using the Wang method and with a p-value filter threshold of 0.05 (Harbig et al. 2023). Inputs were gene lists of the spider and harvestman genes that were in ohnologue or single-copy orthologue gene families, or were early- or late-specific in expression.

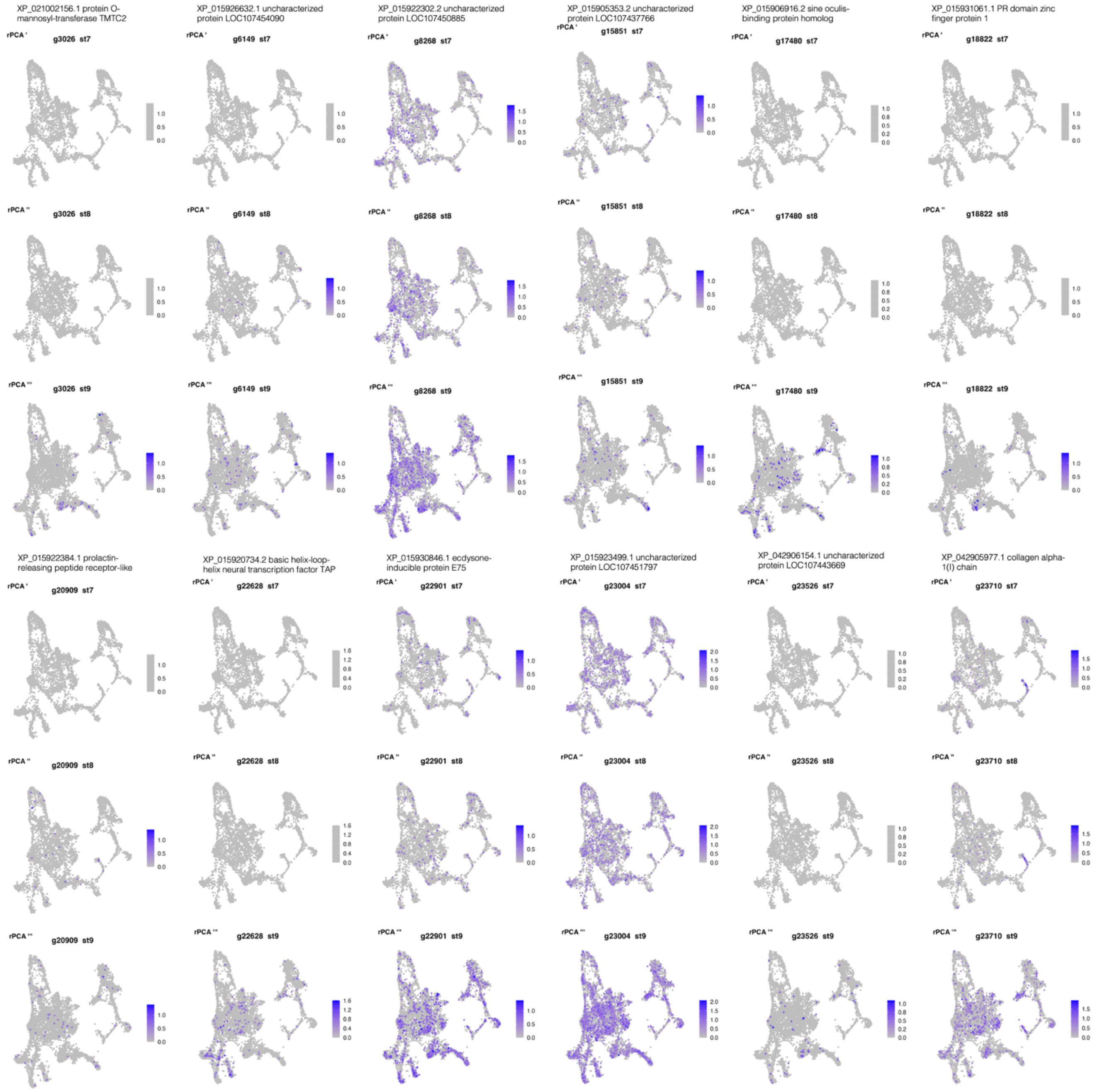


Figure S4: Expression of the rest of the late-specific genes with late-specific peaks in the single cell atlas of stages 7, 8, and 9 for *P. tepidariorum* (Leite et al. 2024). Genes with late-specific expression and peaks but not found in SC data: Pt-g5649, Pt-g12738, Pt-g13508, Pt-g19955.

*Description of Supplementary Files*

Supplementary File 1: S1_PDD_synteny.xlsx

Locations of *Pitx*, *Dmbx*, and *Drgx* genes in arachnids, as described in Aase-Remedios et al. (2023), with the new *P. opilio* annotation included.

Supplementary File 2: S2_PopiV2HDs.xlsx

The full annotation of all homeobox genes with homeodomain sequences in the new *P. opilio* assembly sequenced in this study.

Supplementary File 3: S3_hboxatacprofiles.zip

To-scale figures of ATAC profiles from each stage of conserved homeobox gene clusters. Filenames represent the species, the genes included, and the gene IDs for that sub-cluster, relative to Figure 7a.
