## Supplementary File 3 for "Assessing the impact of whole genome duplication on gene expression and regulation during arachnid development": READ_ME.txt

Information to help interpret the ATAC profiles of conserved homeobox gene clustering.

Figures are plotted to scale.

Filenames indicate the species, cluster/sub-cluster and the gene ID #
The gene IDs are the leftmost and rightmost genes in that cluster/sub-cluster.

Figures contain intervening non-homeobox genes.

Species:
po: Phalangium opilio (harvestman)
pt: Parasteatoda tepidariorum (spider)


Hox:
The Hox cluster has been subdivided into three sub-clusters:
ant (anterior): lab - pb - Hox3
mid (middle): Dfd - Scr - ftz
post (posterior): Antp - Ubx - abd-A - Abd-B

NK subclusters:
spider NK-core (NK7-A, Lbx-A, Tlx-A, NK3, and NK4)
po NK7 - Lbx and po Tlx - NK3
and spider Lbx-B - Tlx-B
Emx1 - Emx2 in all species
and po NK5 - NK1 and spider NK1-B-2 - NK5

Irx: all genes as depicted in the figure

HRO:
Otp - Hbn - Rax-A in spider
Otp - Rax  - Hbn in harvestman
Isl or spider Rax-B not included

SINE
Po: as depicted in the figure
Pt-A Six4/5-A - Six1/2-A
Pt-B as depicted in the figure
