## Supplementary figures and images for "Assessing the impact of whole genome duplication on gene expression and regulation during arachnid development"

### po_emx1_emx2_pog21490-pog21492.png

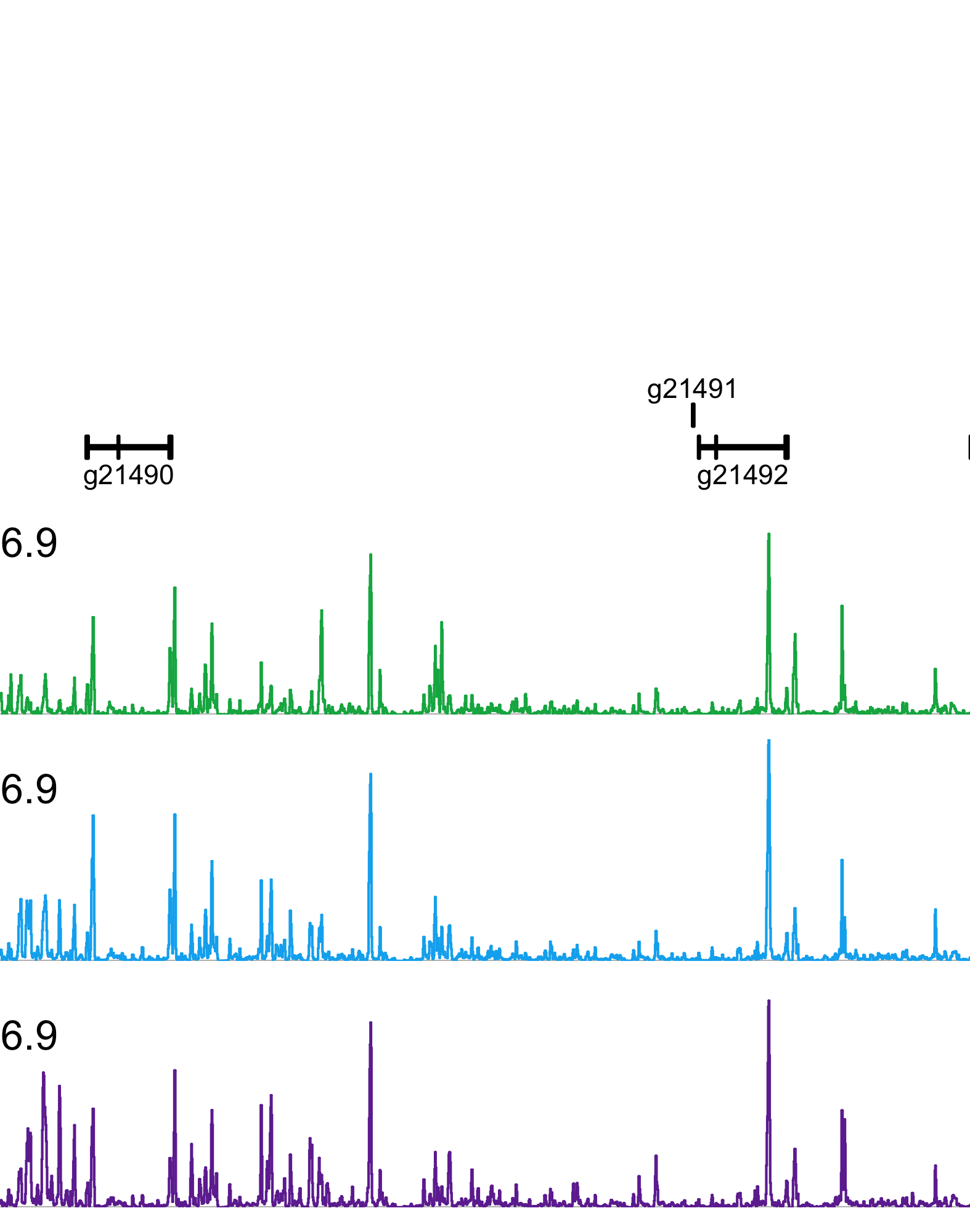

### po_hox_ant_pog6952-pog6956.png

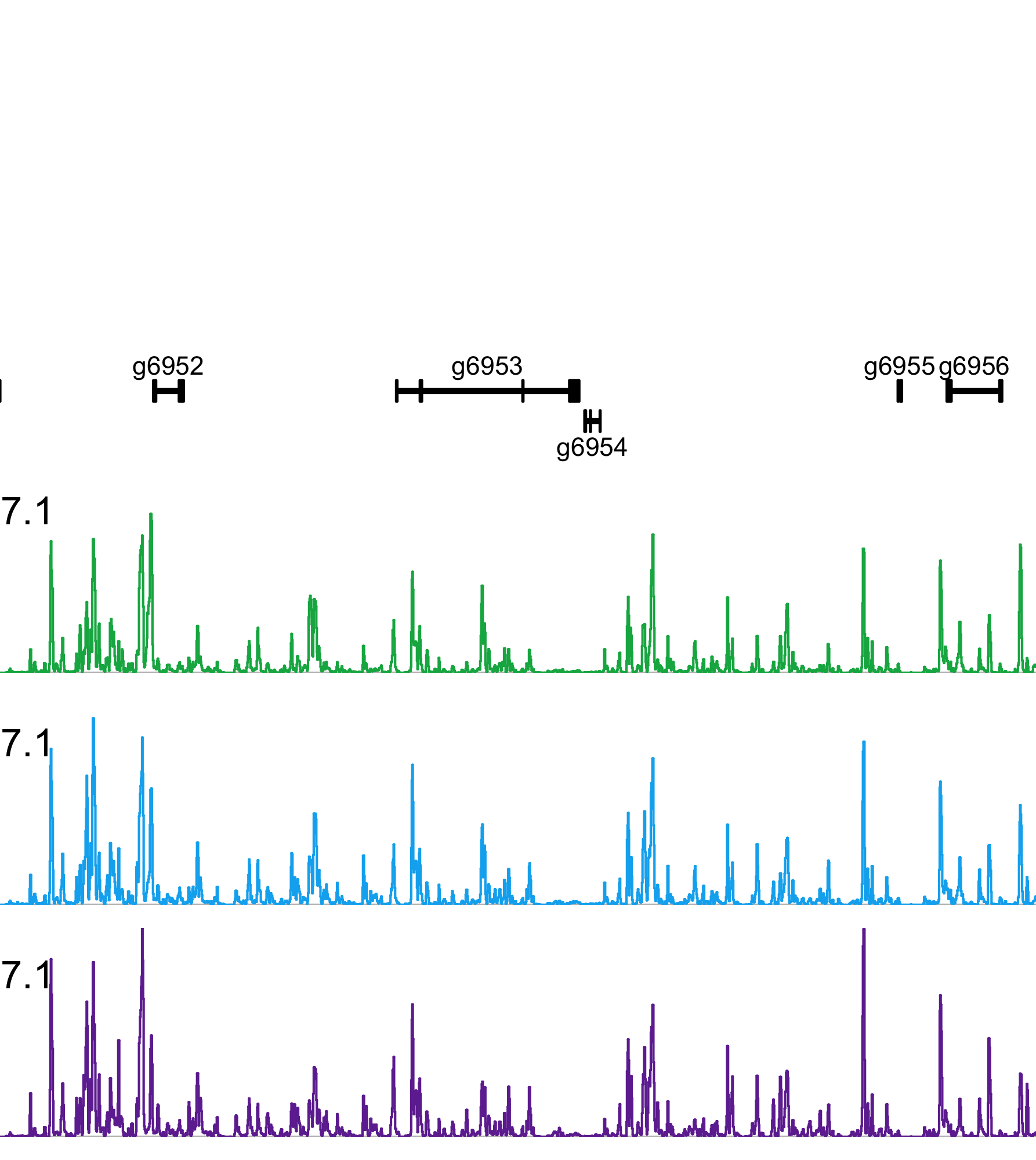

### po_hox_mid_pog6945-pog6947.png

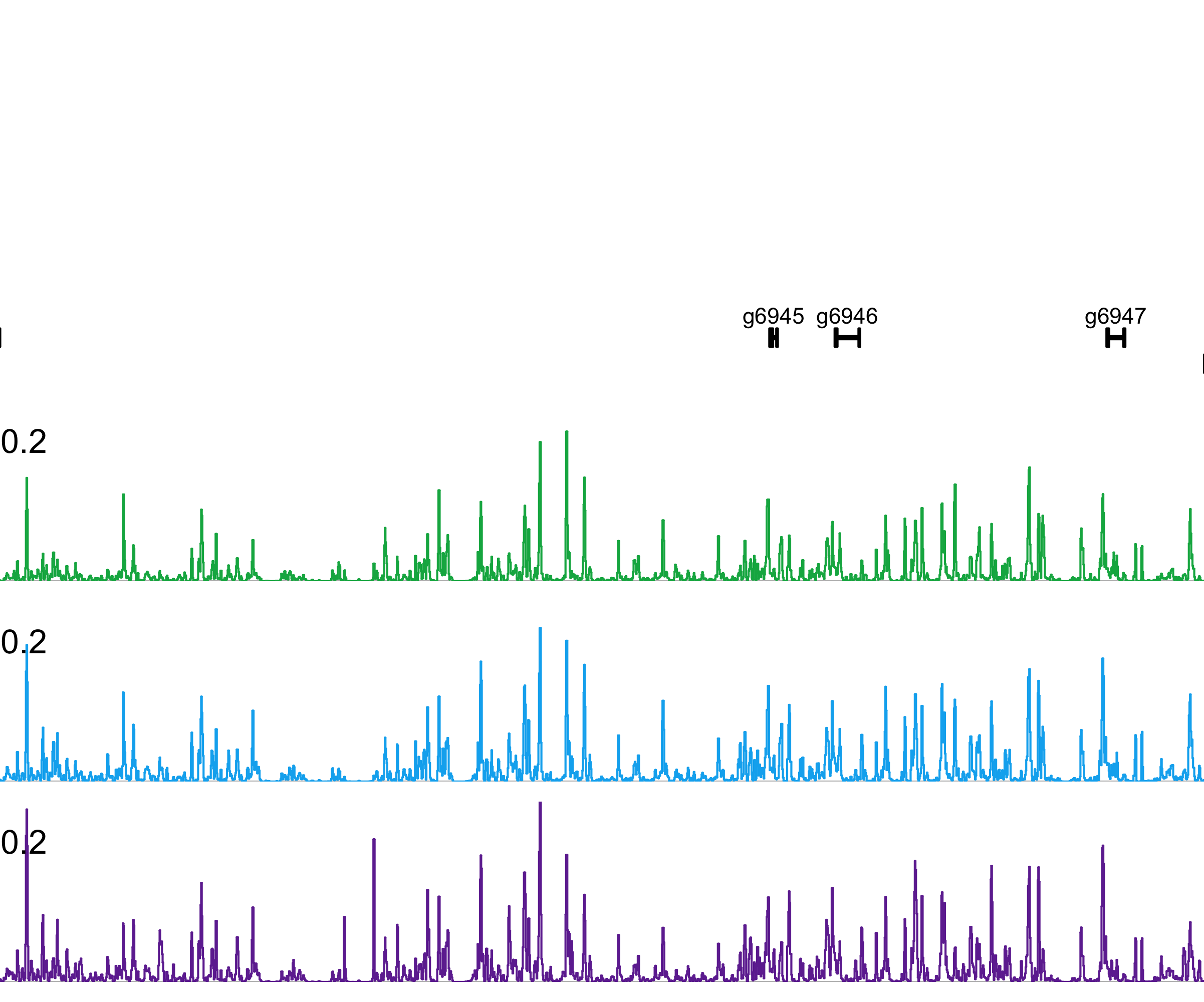

### po_hox_post_pog6938-pog6944.png

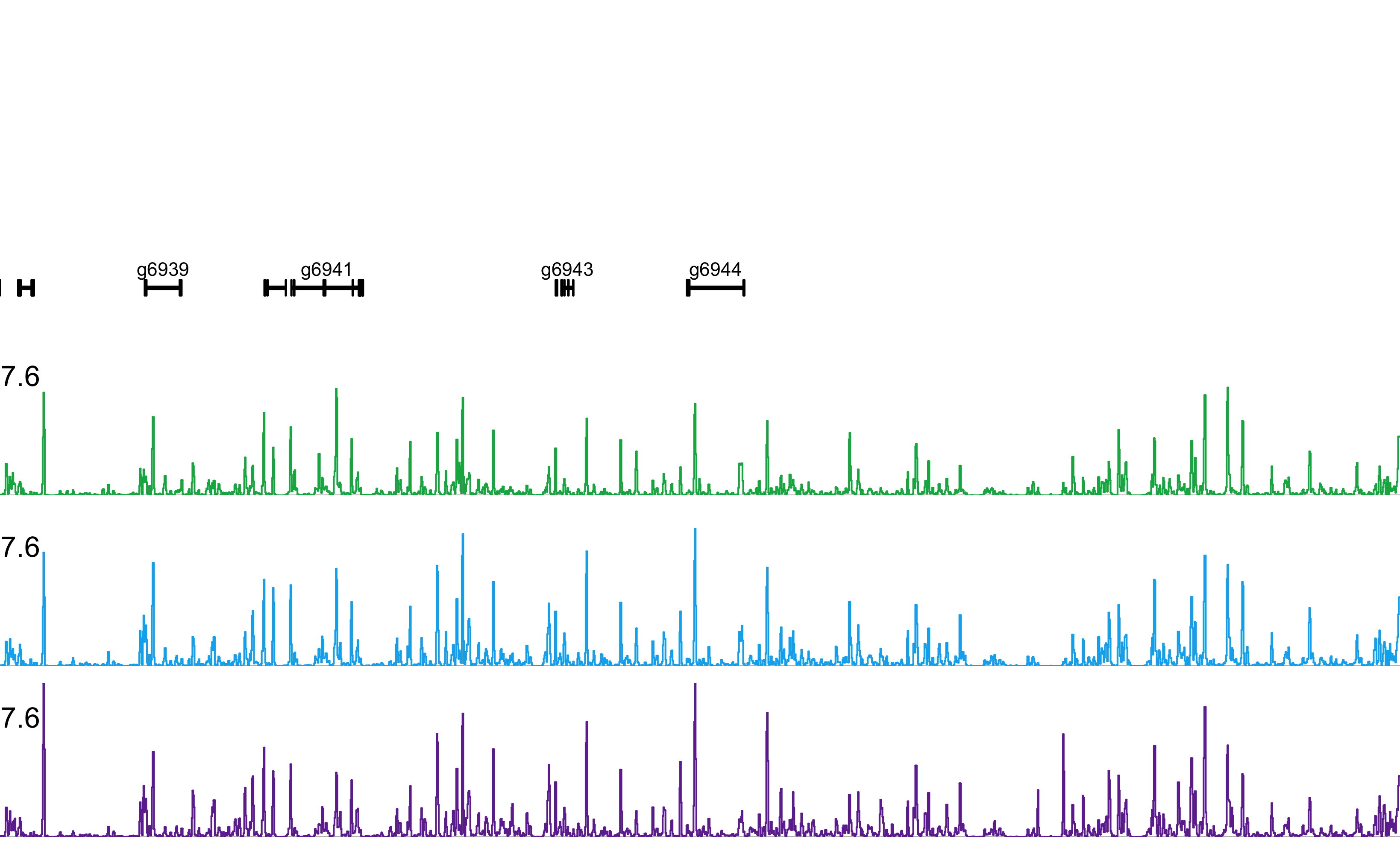

### po_hro_pog6912-pog6914.png

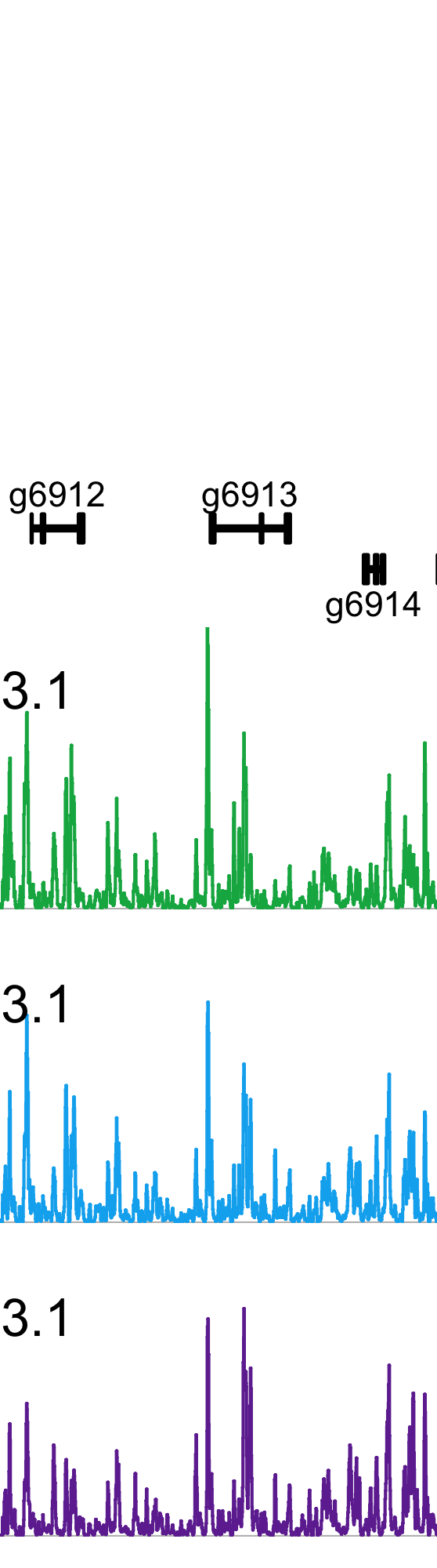

### po_irx_pog15350-pog15353.png

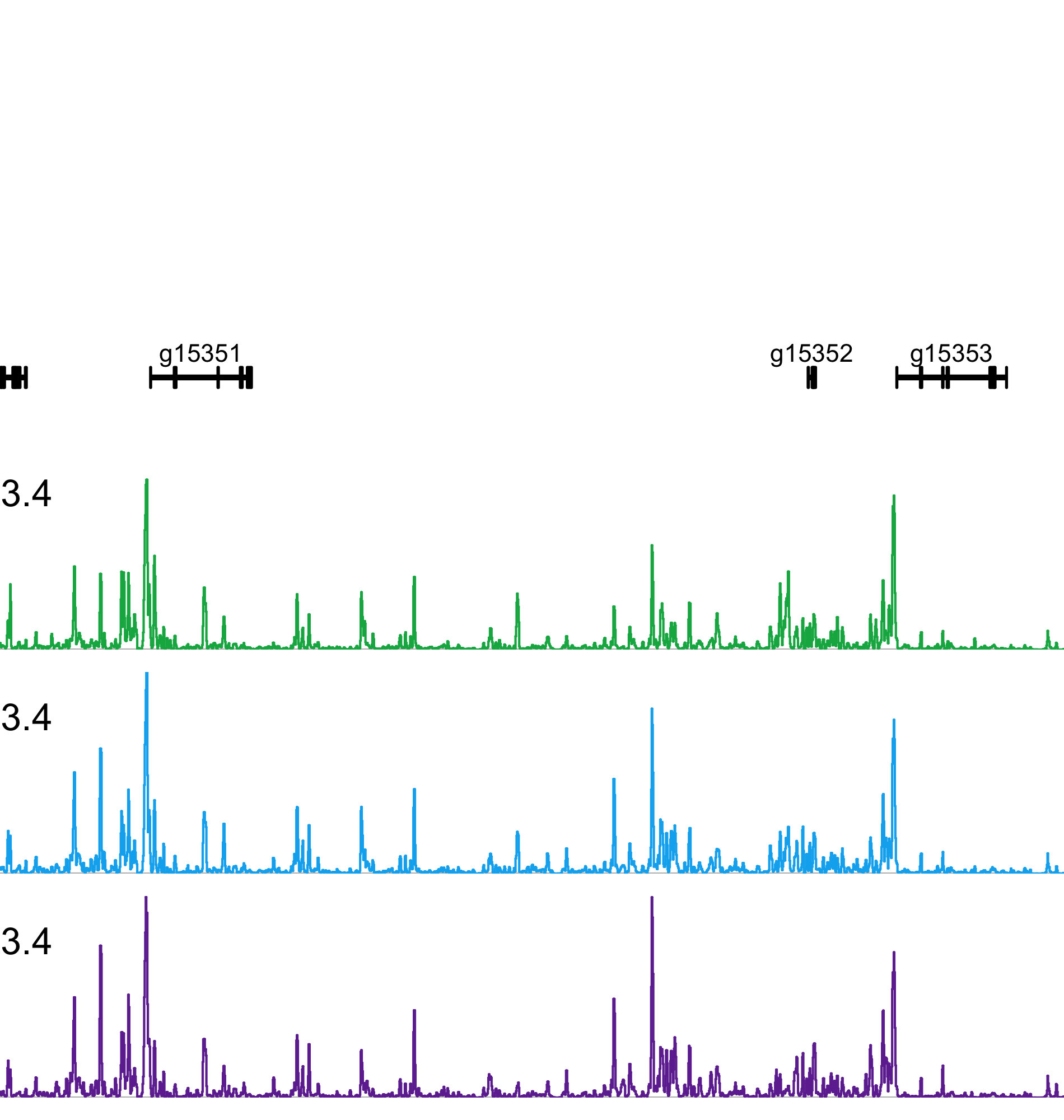

### po_nk5_nk1_pog21726-pog21728.png

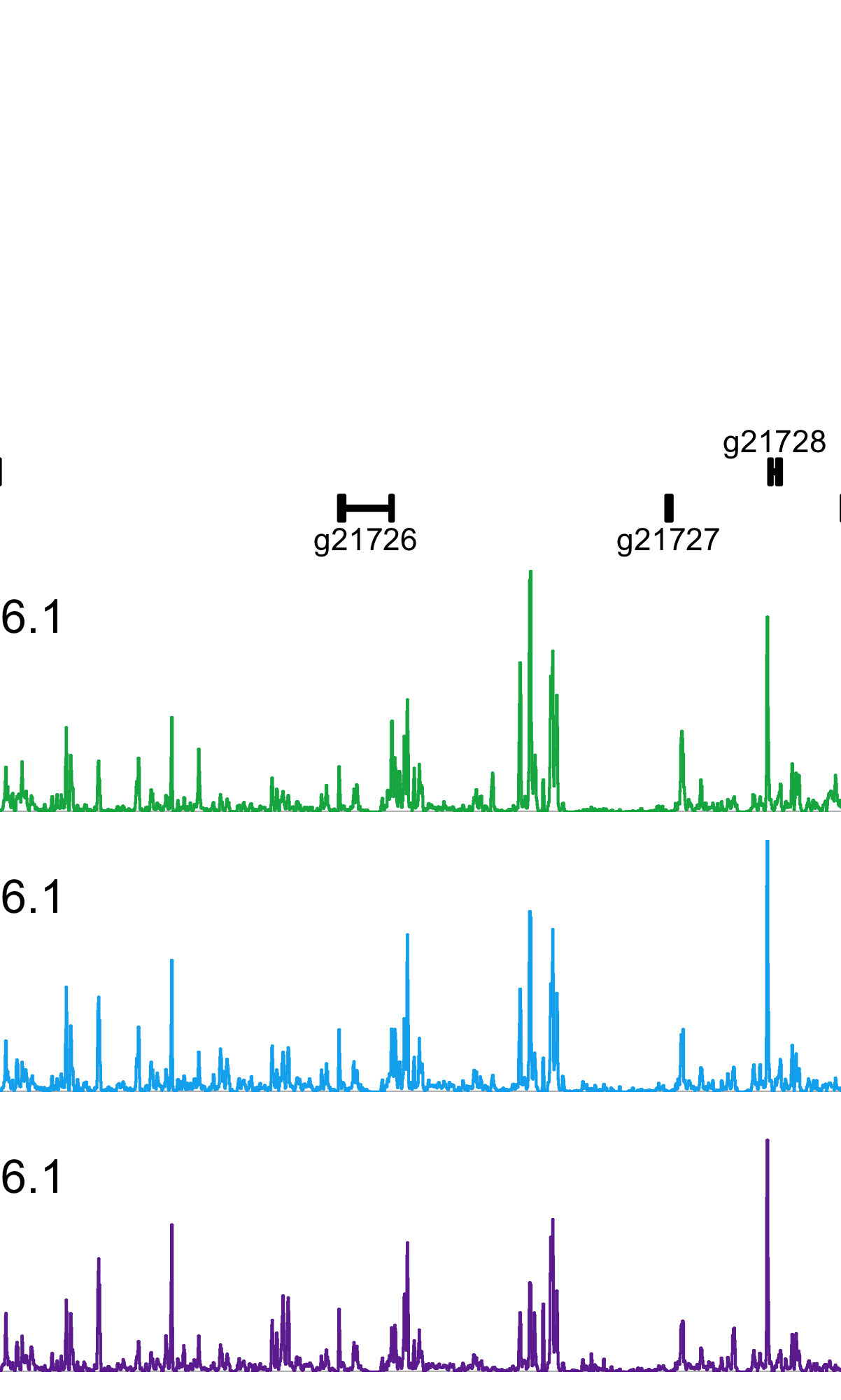

### po_nk7_lbx_pog12032-pog12036.png

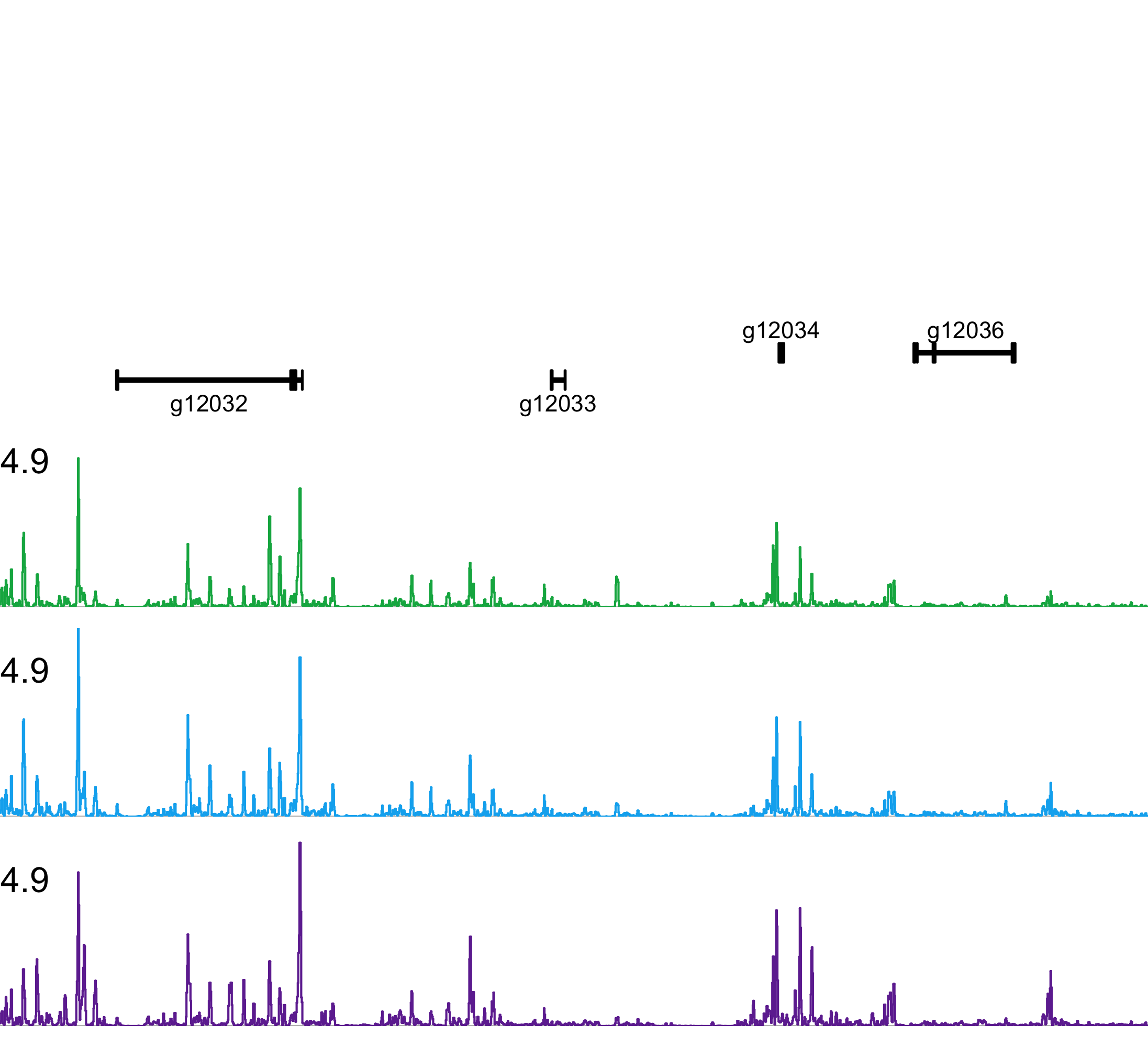

### po_six36_six45_pog14172-pog14182.png

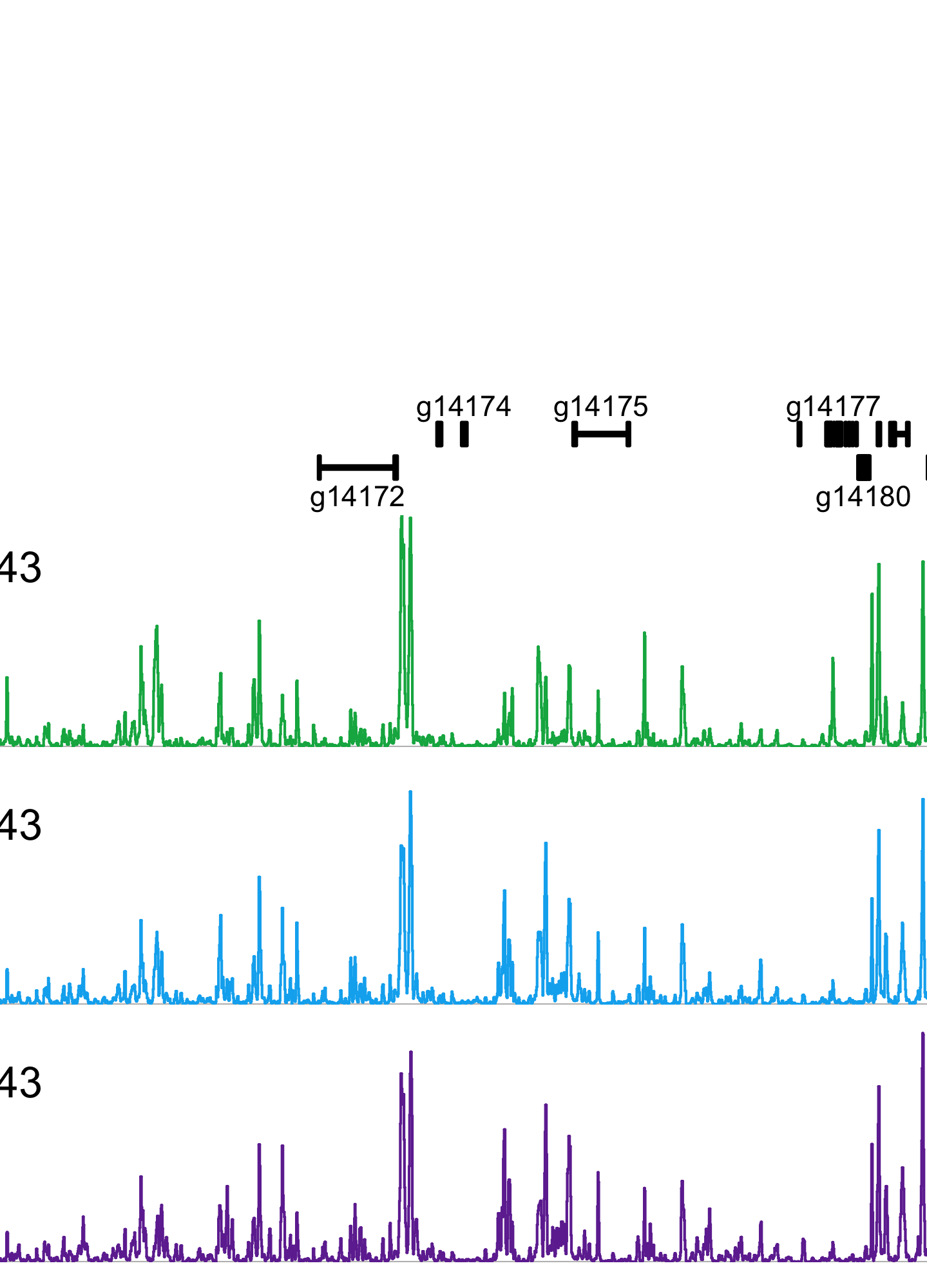

### po_tlx_nk3_pog21708-pog21710.png

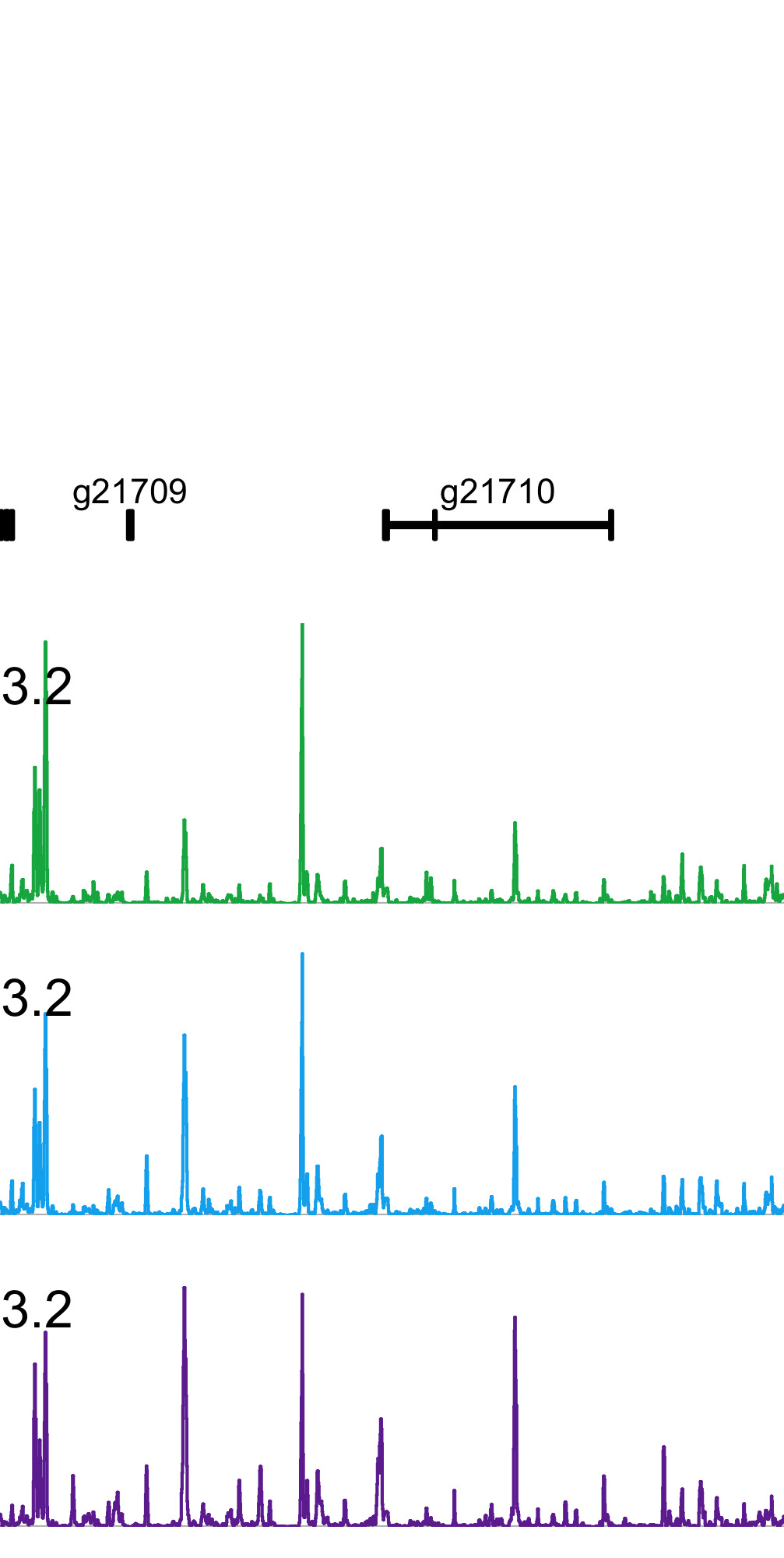

### pt_hro_ptg16269-ptg16274.png

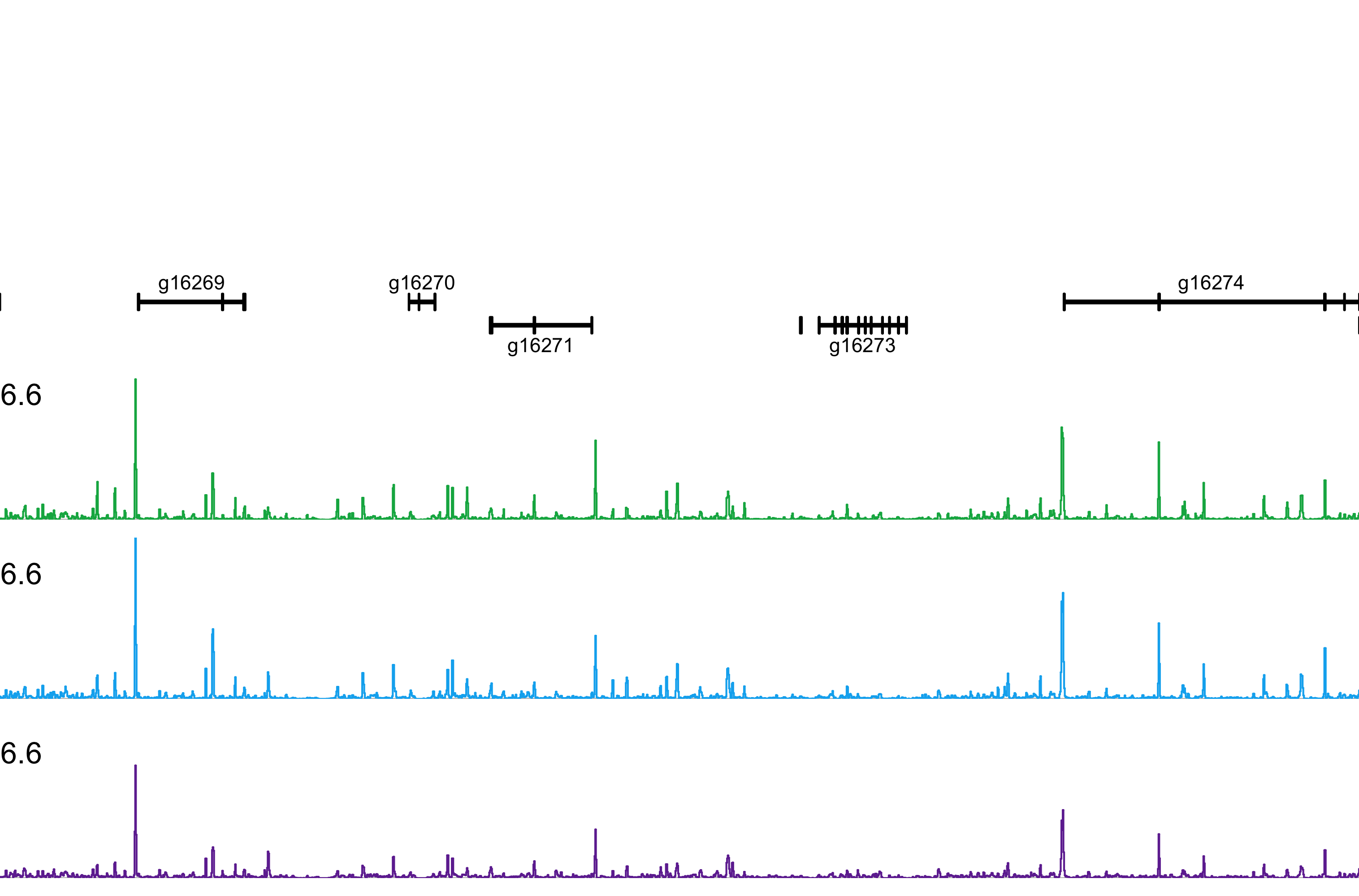

### pt_nk5_nk12_ptg2489-ptg2490.png

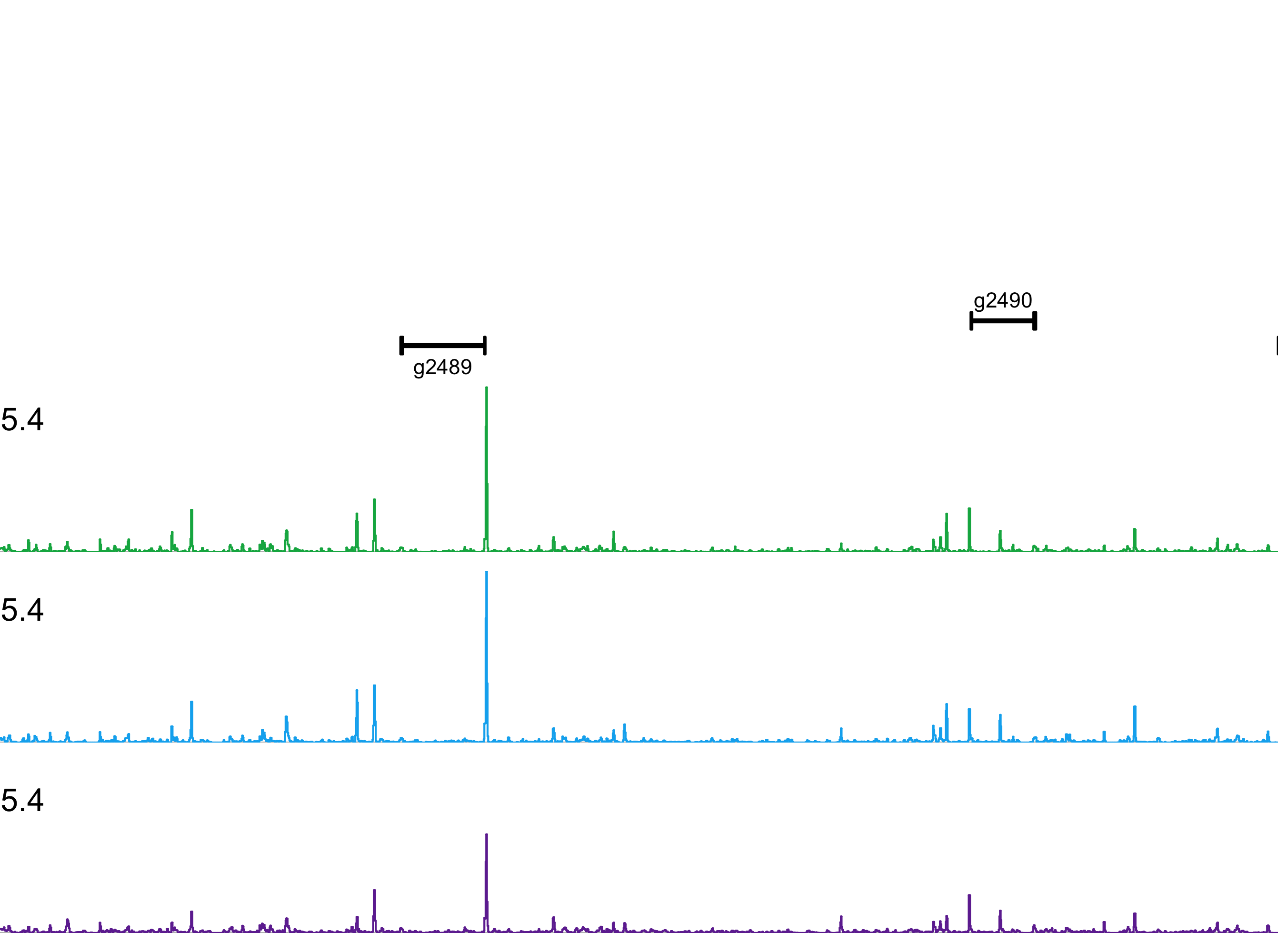

### ptD_emx1_emx2_ptg2828-ptg2829.png

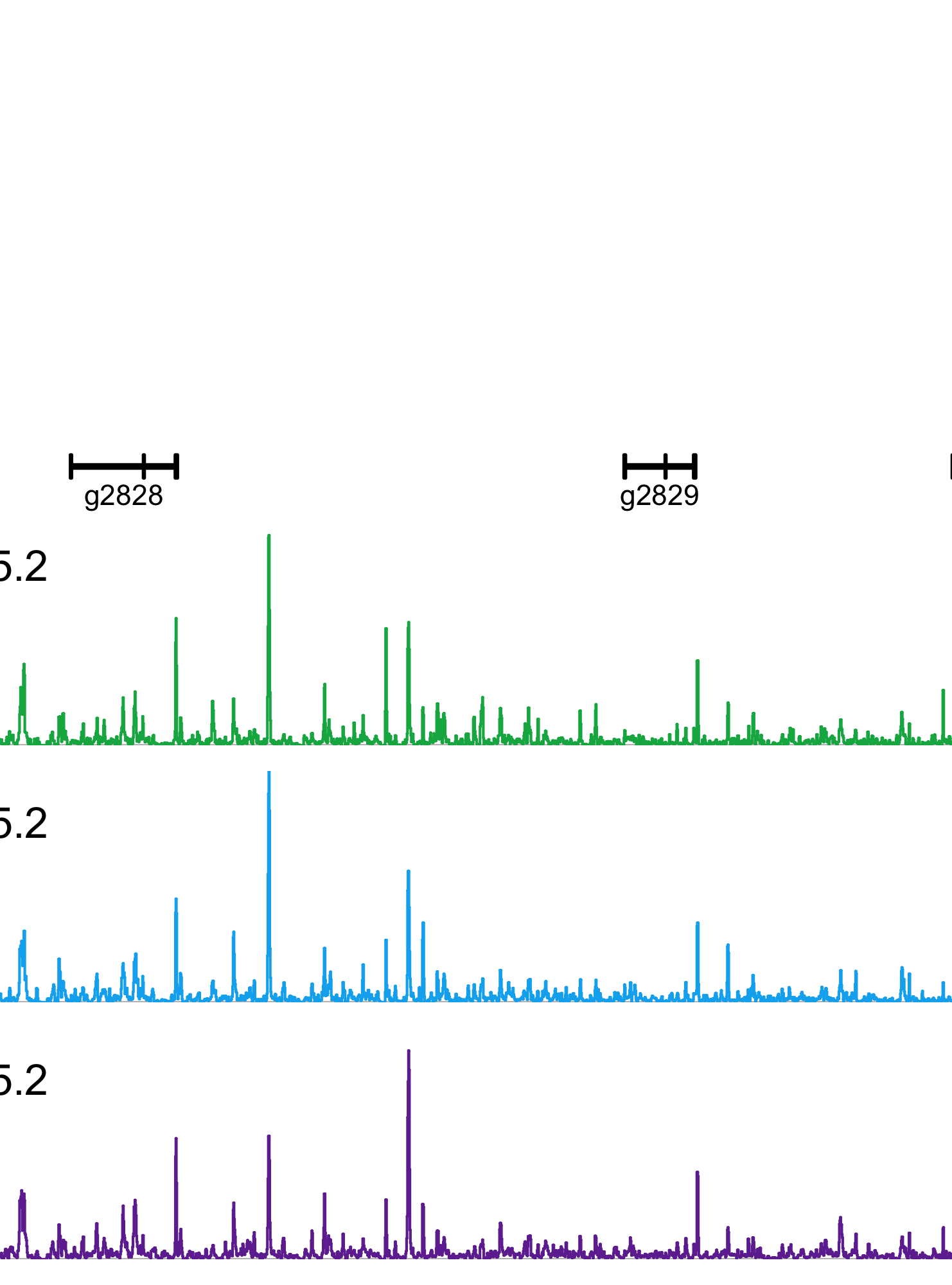

### ptD_hox_ant_ptg11723-ptg11725.png

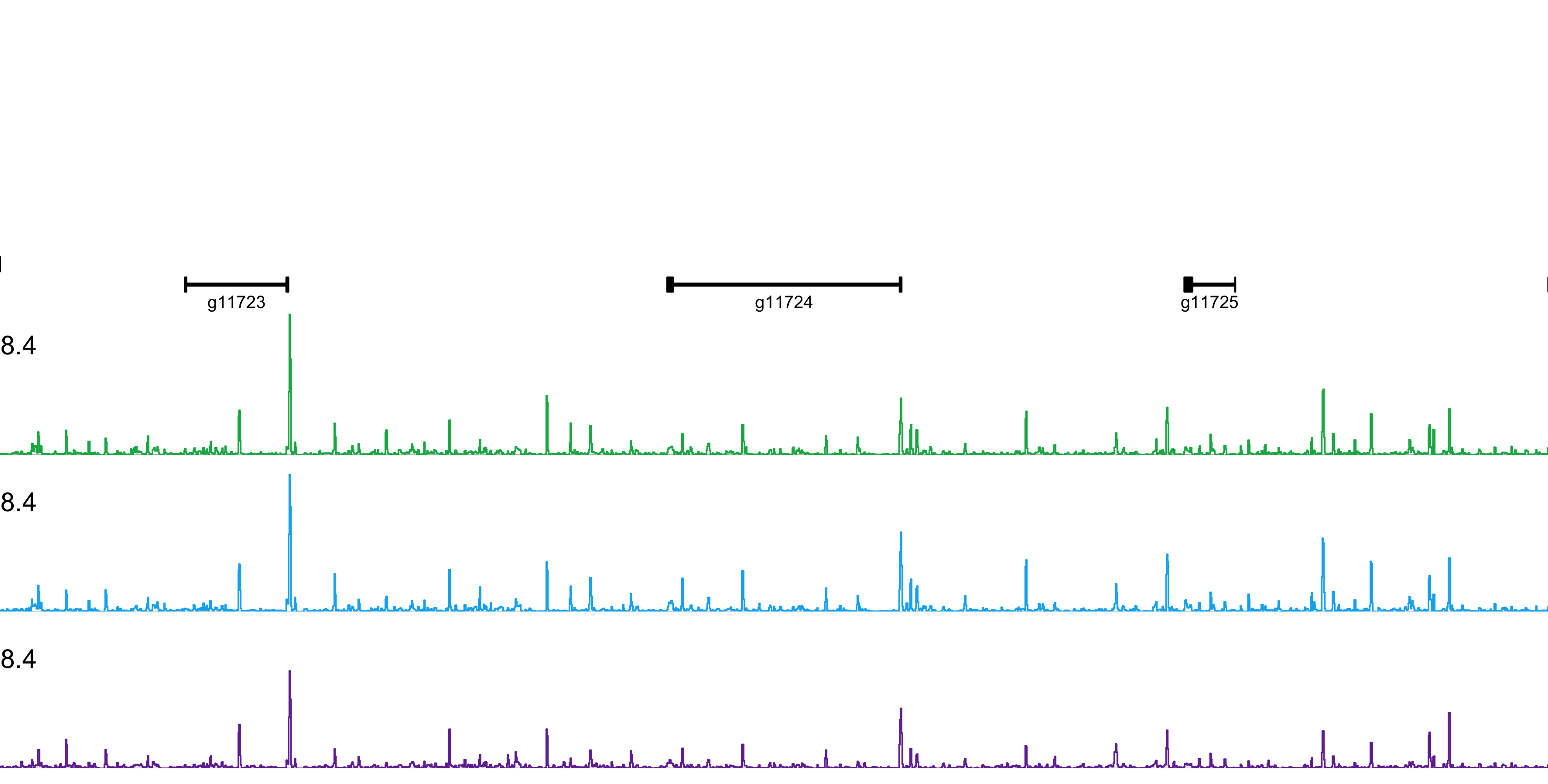

### ptD_hox_mid_ptg11749-ptg11753.png

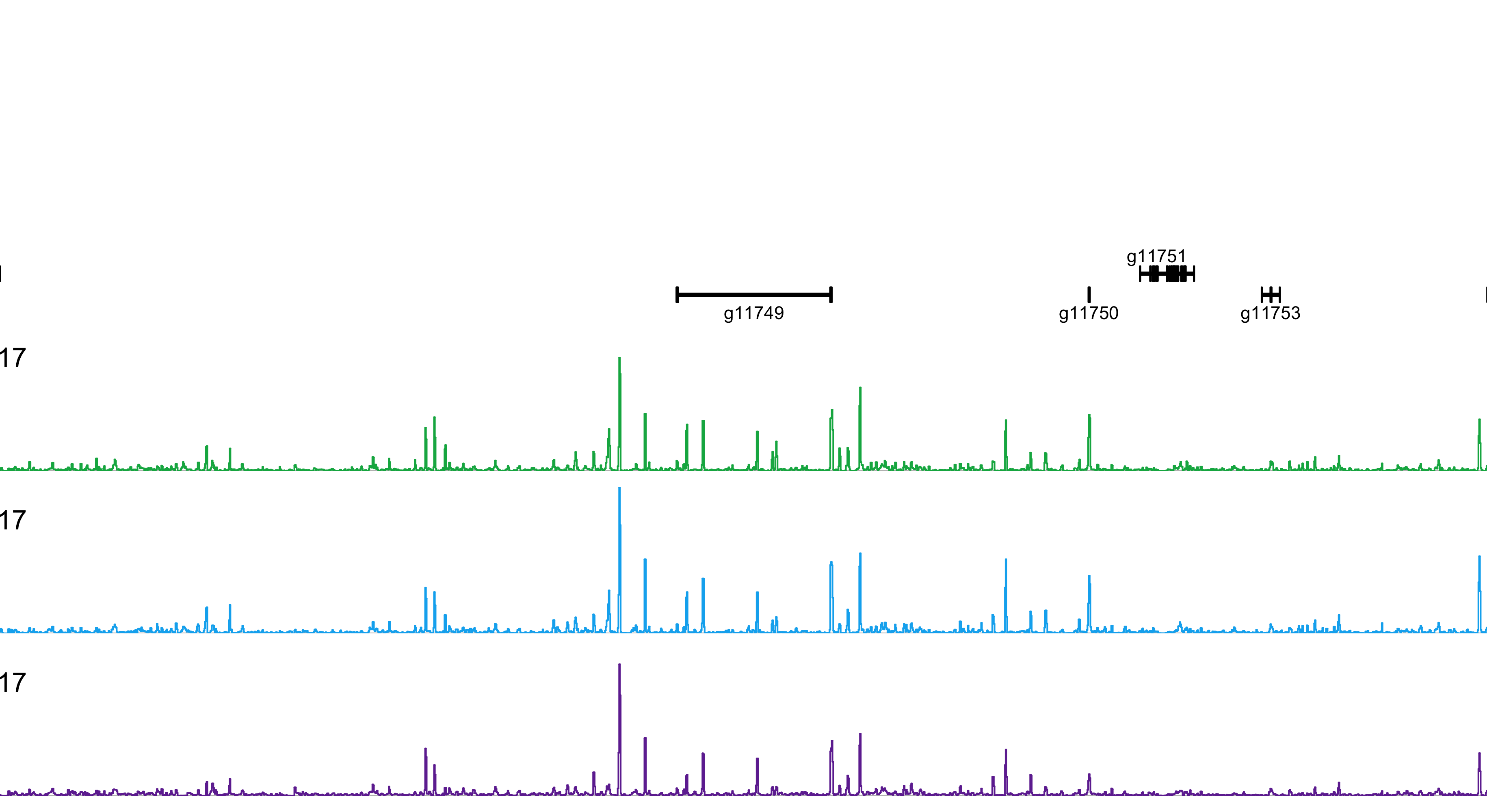

### ptD_hox_post_ptg11761-ptg11768.png

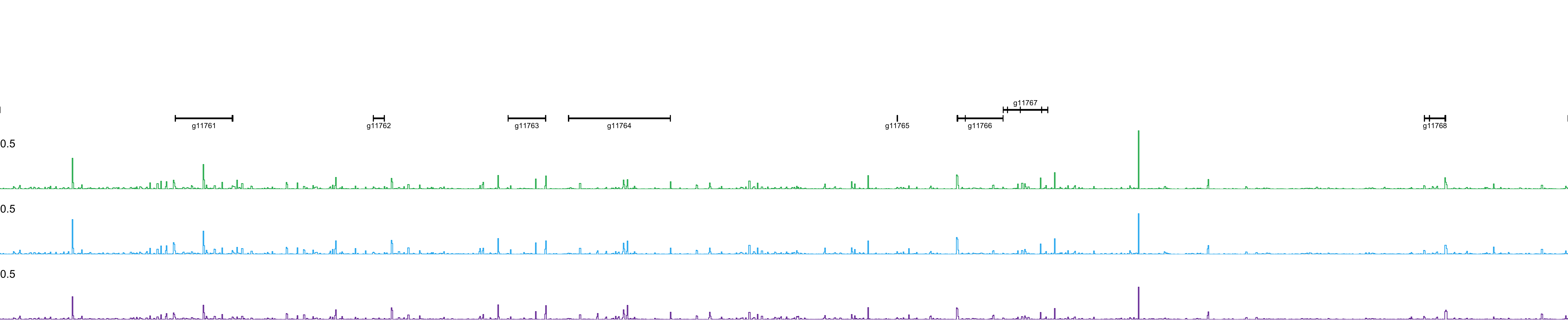

### ptD_irx_ptg23939-ptg23942.png

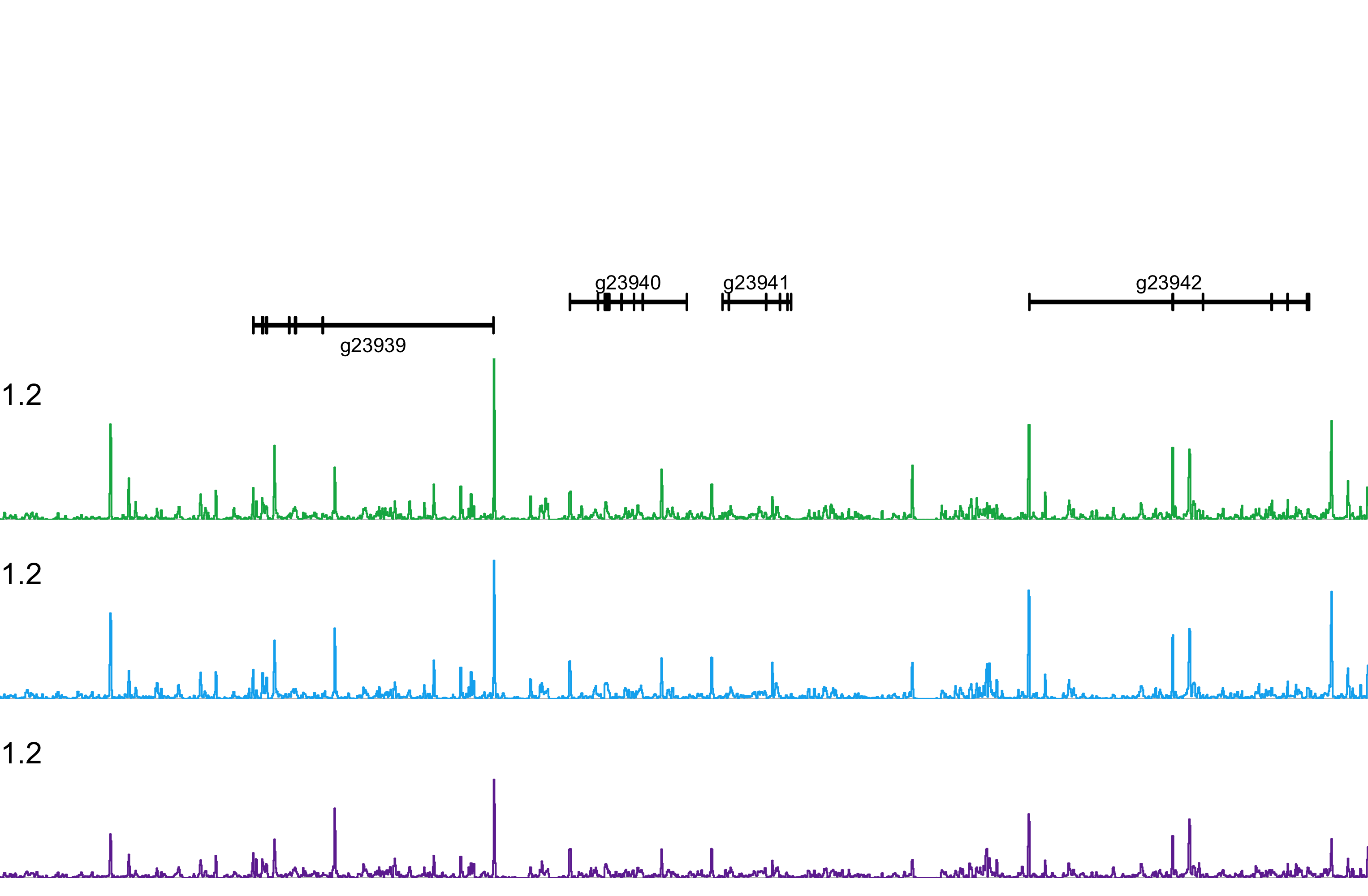

### ptD_lbx_tlx_ptg2193-ptg2195.png

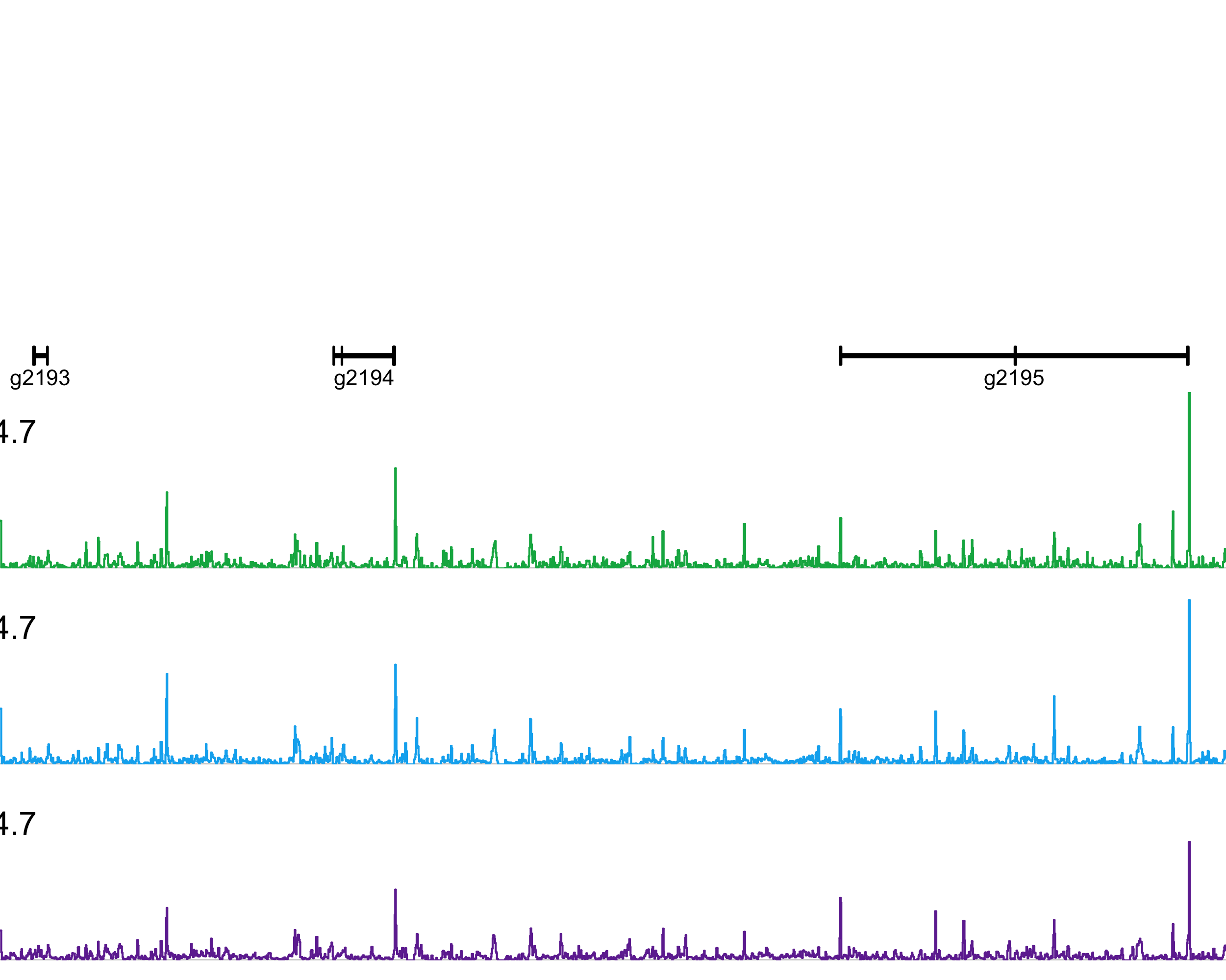

### ptD_six36_six12_ptg4657-ptg4659.png

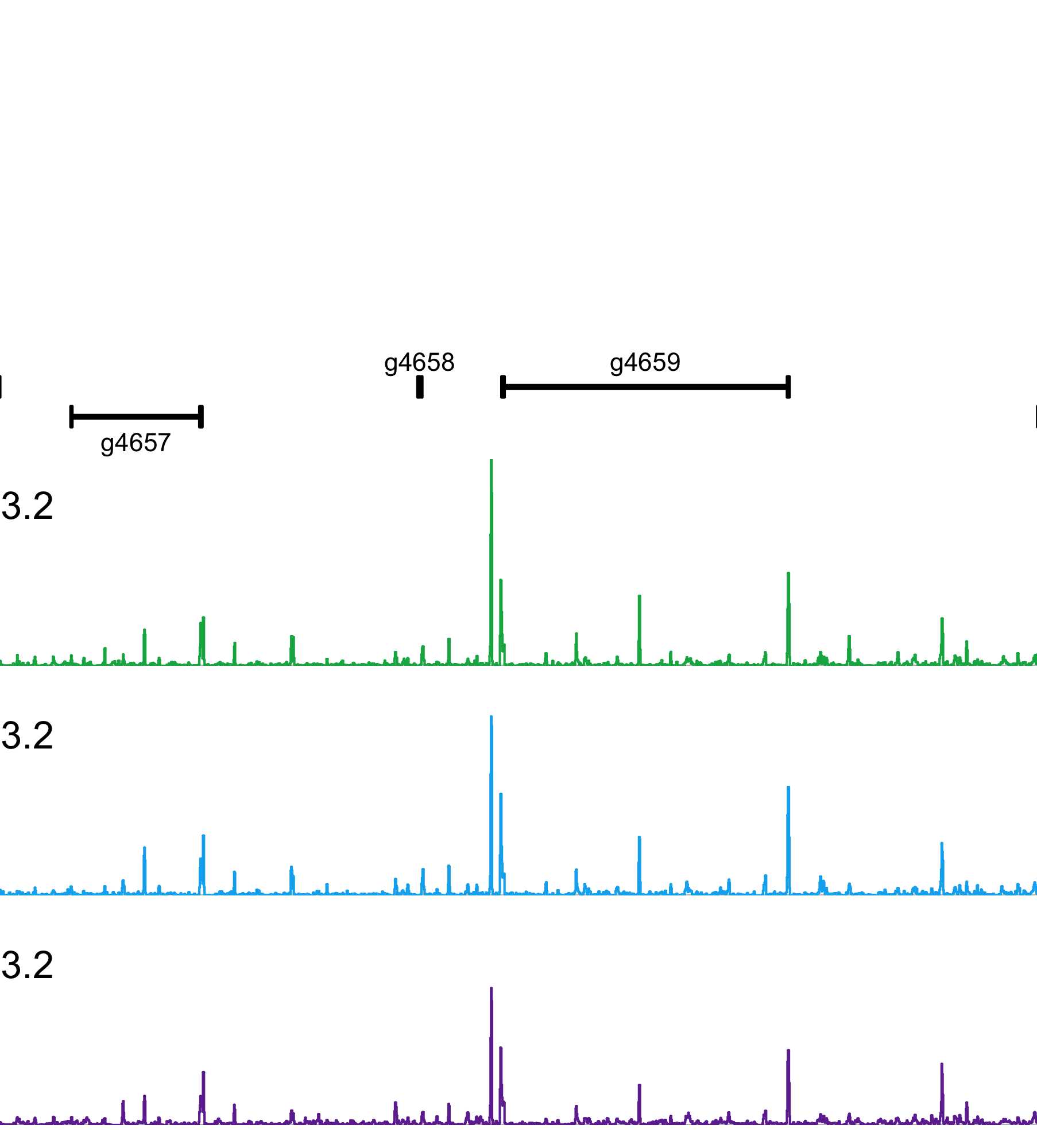

### ptI_emx1_emx2_ptg9266-ptg9267.png

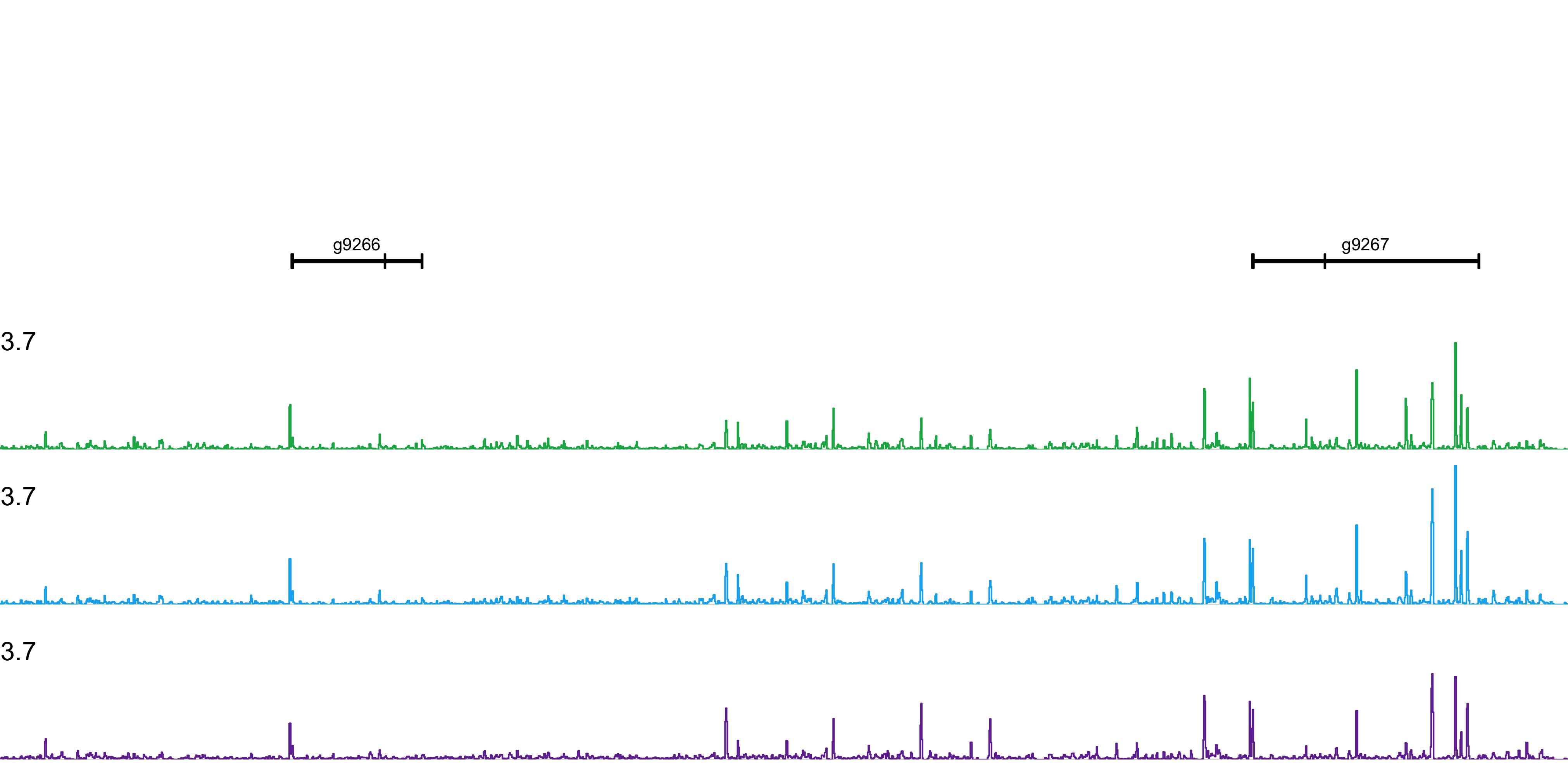

### ptI_hox_ant_ptg3430-ptg3438.png

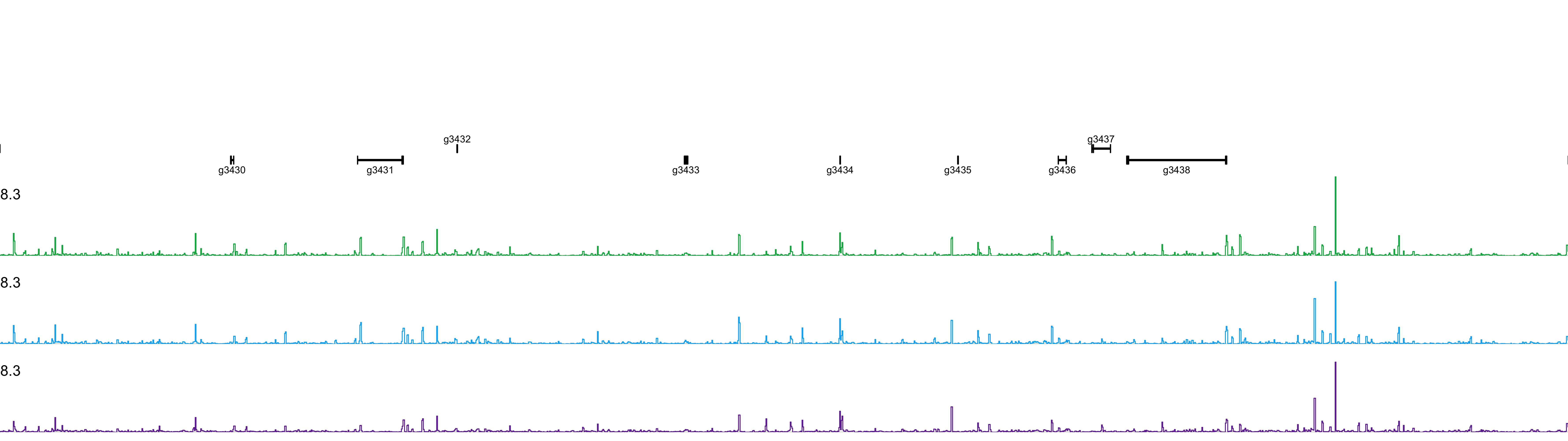

### ptI_hox_mid_ptg3449-ptg3452.png

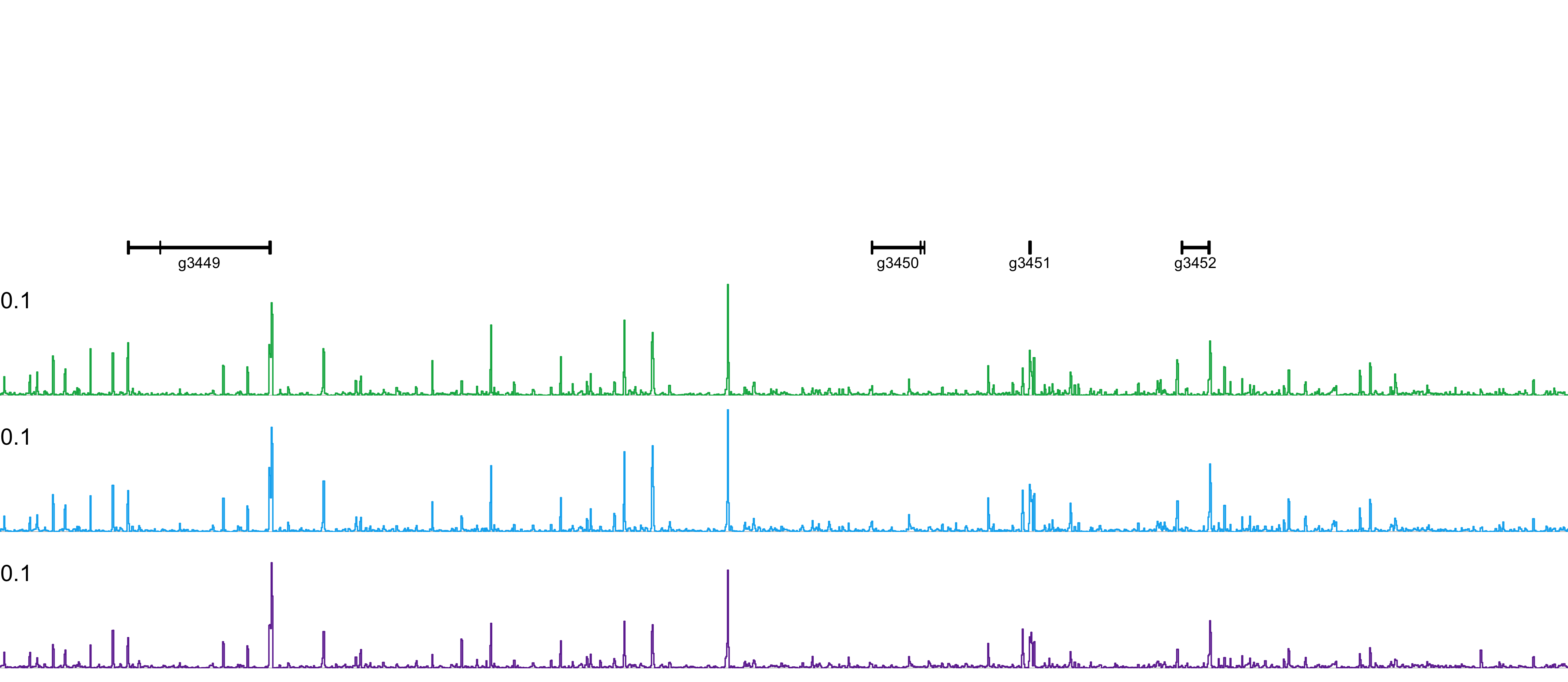

### ptI_hox_post_ptg3457-ptg3466.png

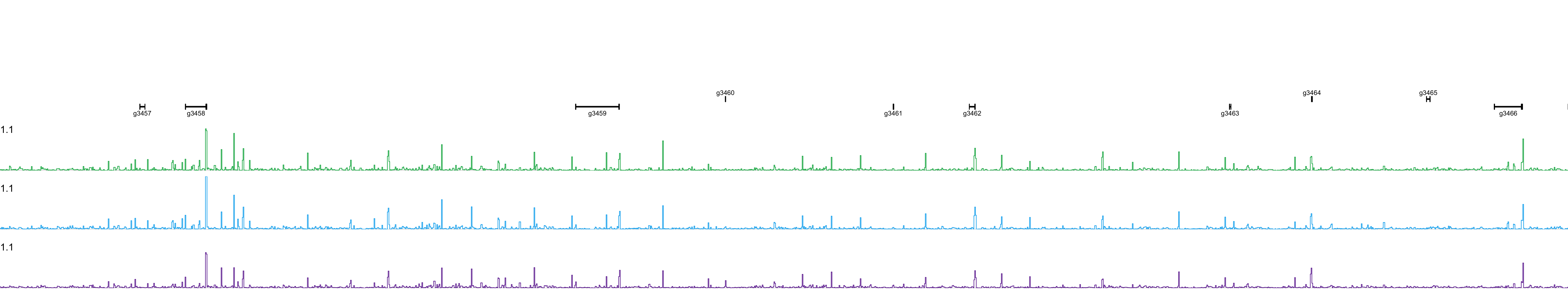

### ptI_irx_ptg15773-ptg15778.png

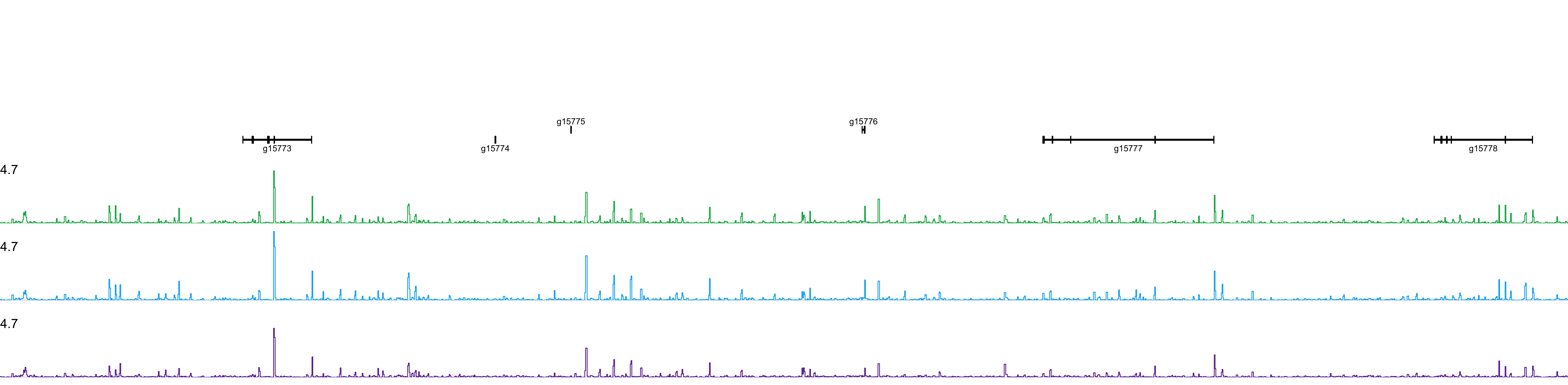

### ptI_nk_core_ptg21439-ptg21445.png

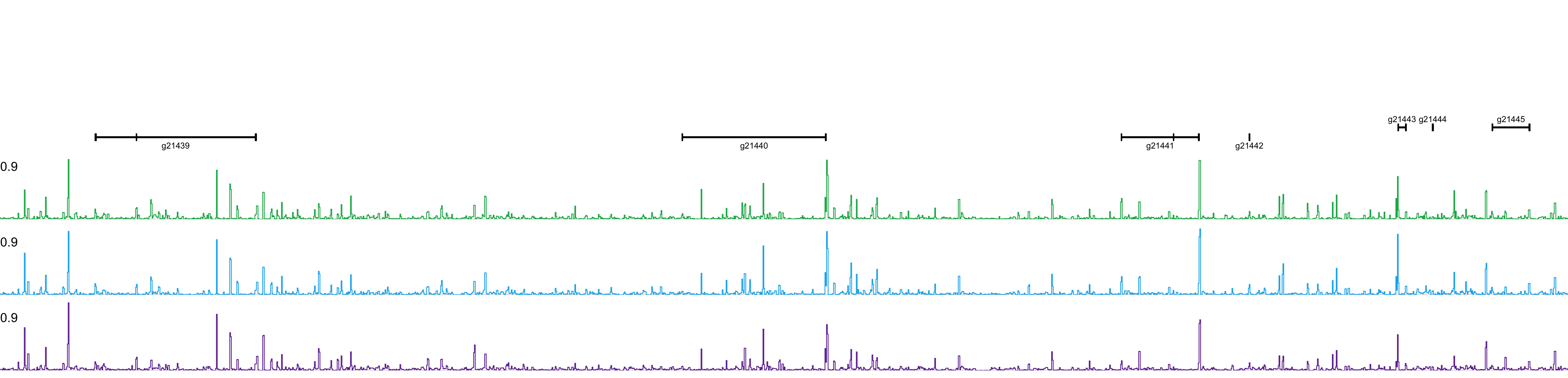

### ptI_six12_six45_ptg11900-ptg11902.png

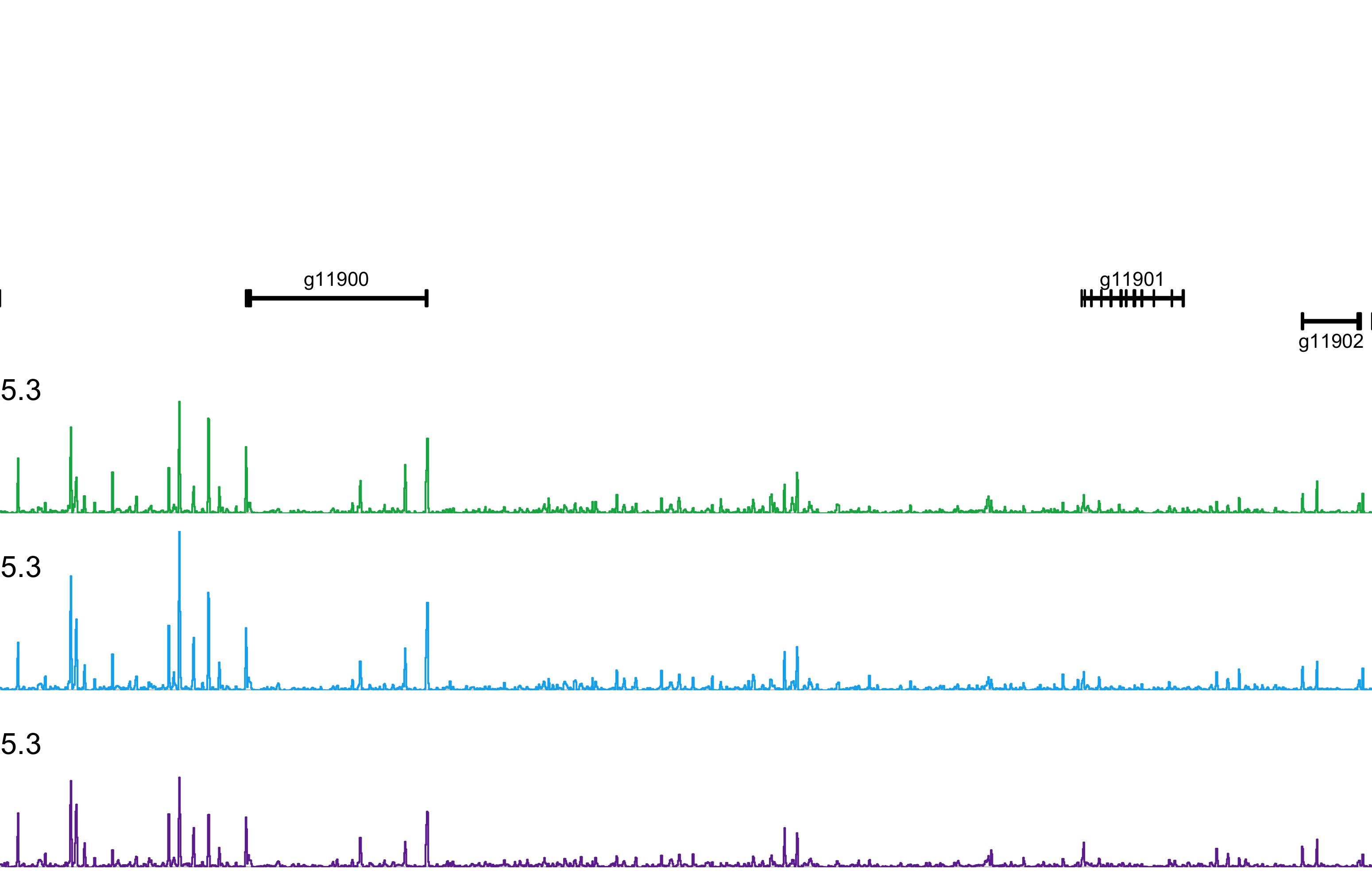
